# Dysfunctional analysis of the pre-training model on nucleotide sequences and the evaluation of different k-mer embeddings

**DOI:** 10.1101/2022.12.05.518770

**Authors:** Yao-zhong Zhang, Zeheng Bai, Seiya Imoto

## Abstract

Pre-training has attracted much attention in recent years. Although significant performance improvements have been achieved in many downstream tasks using pre-training, the mechanism of how a pre-training method works for downstream tasks is not fully illustrated. In this work, focusing on nucleotide sequences, we decompose a pre-training model of Bidirectional Encoder Representations from Transformers (BERT) into embedding and encoding modules to illustrate what a pre-trained model learns from pre-training data. Through dysfunctional analysis on both data and model level, we demonstrate that the context-consistent k-mer representation is the primary product that a typical BERT model learns in the embedding layer. Surprisingly, single usage of the k-mer embedding pre-trained on the random data can achieve comparable performance to that of the k-mer embedding pre-trained on actual biological sequences. We further compare the learned k-mer embeddings with other commonly used k-mer representations in downstream tasks of sequence-based functional predictions and propose a novel solution to accelerate the pre-training process.

**Supplementary information:** The source code and relevant data are available at https://github.com/yaozhong/bert_investigation.

## 1 Introduction

Pre-training using a transformer-based model has been achieving ground-breaking results in many research fields, such as natural language processing and computer vision (Devlin *et al*., 2018; Dosovitskiy *et al*., 2020). In computational biology, the technique has been applied to learn representations of amino acid sequences and nucleotide sequences (Rao *et al*., 2021; Jumper *et al*., 2021; Ji *et al*., 2021). As the amount of labeled data is very few, the pre-training and fine-tuning paradigm leverages available unlabeled data to learn a general representation and further refines learned model parameters using limited labeled data from a downstream task. Although significant performance improvements have been achieved in many downstream tasks using such a training paradigm, the mechanism of how a pre-trained model works is not fully illustrated. Moreover, most current works directly use the model architecture that is the same as in natural language modeling tasks, while biological-sequence-specific characters are less explored.

In this work, focusing on nucleotide sequences, we aim to interpret what a pre-trained Bidirectional Encoder Representations from Transformers (BERT) model (Devlin *et al*., 2018; Ji *et al*., 2021) learns through pre-training and how it works in downstream tasks. In the previous work (Ji *et al*., 2021), attention weights of the model are used to interpret important regions contributing to the prediction for a specific input sequence. However, non-sequence-specific token embeddings learned by the pre-trained model are yet to be fully explored. Here, we focus on the non-sequence-specific lower-level token embeddings to give a global interpretation (Novakovsky *et al*., 2022) of the pre-trained model. We decompose the BERT model into the embedding and encoding modules. We use a dysfunctional approach to investigate different modules by incorporating randomness on both data and model levels. Besides evaluating different pre-trained models, we also analyze the use of different k-mer embeddings in the sequence-based functional prediction tasks of TATA promoter prediction and transcript factor binding site (TFBS) prediction. Finally, we propose a potential usage of the BERT model pre-trained on randomly generated sequences to accelerate the pre-training process.

## 2 Methods

To investigate what a BERT model learns through pre-training, we decomposed a typical BERT model into modules of embedding and encoding. The embedding module consists of different embeddings representing tokens, token types, and positions. The encoding module is the encoder part of a transformer (Vaswani *et al*., 2017), which contains multiple attention and feed-forward layers. For analyzing the pre-training of the BERT model on nucleotide sequences, we made use of DNABERT (Ji *et al*., 2021). ^1^

In the pre-training, partial input tokens are masked, and the model is trained to predict these masked tokens. The masking strategy used in DNABERT is illustrated in Figure 1. To avoid a masked token being trivially inferred through the immediately surrounding k-mers, DNABERT masks k contiguous k-mers. For example, the five contiguous 5-mers are masked for the pre-training, as shown in Figure 1. If a k-mer token is further decomposed into nucleotide level, **we can find that partial nucleotides in the masked k-mer can still be inferred from their immediately surrounding k-mers. As a result, for k contiguous masked k-mers, only one nucleotide can not be trivially determined from their surrounding k-mers**. For the DNABERT model, the label search space is (4^*k*^ + 5) (five labels are special tokens as used in the BERT(e.g., [CLS], [PAD])), while the non-inferable label search space is 4 + 5. Therefore, the pre-training task used in DNABERT can be decomposed into two sub-tasks: (1) context-consistent k-mer prediction for the contiguous masked k-mers. (2) predicting the non-trivial-inferable nucleotide in k-mers. For different pre-training data, such as nucleotide sequences of different species, the first sub-task is species-independent, while the second sub-task is data-dependent. According to the previous work of predicting nucleotide from its surrounding nucleotide (Liang *et al*., 2022), the performance is different for different regions across all human chromosomes. The repeat region has a higher accuracy, while the coding region has a lower accuracy. The overall accuracy is over 50%. This performance correlates with the upper bound of the pre-training. Note that the encoding module takes up a large proportion of model parameters. Taking 5-mer DNABERT for instance, the total number of the model parameter is 86,833,154, while the number of model parameters of the encoding layers is around 98% (85,054,464) of the total ones. For the pre-training on nucleotide sequences, the large model parameters could be trained to make the context-consistent k-mer predictions in the large label search space.

**Fig. 1:**
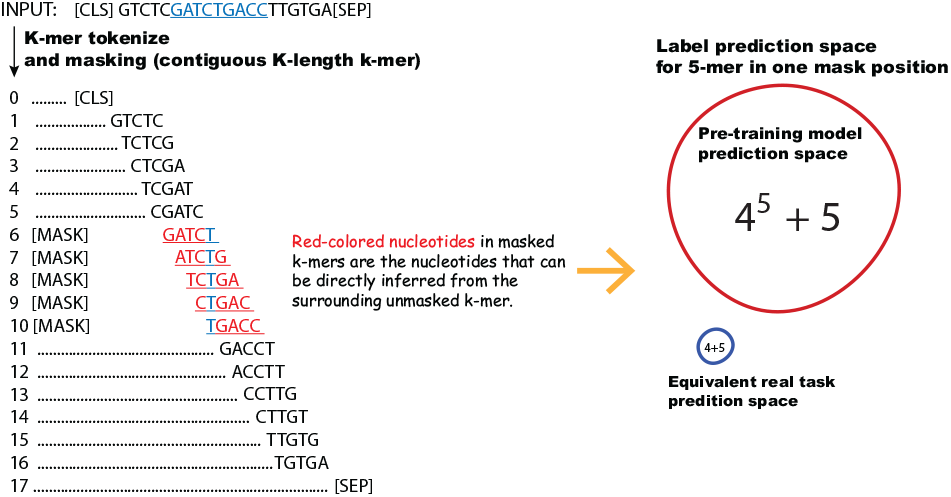
The k-mer sequence masking strategy used in DNABERT. To avoid a masked token from being trivially inferred through the immediately surrounding k-mers, DNABERT masks k contiguous k-mers. Compared with the non-inferable nucleotides predicting sub-task, the original prediction space is huge.

To validate this assumption, **we used a dysfunctional approach to investigate the DNABERT model by incorporating randomness on both data and model levels**. On the data level, we prepared totally randomly generated nucleotide sequences for around 3 billion lengths to compare with the DNABERT pre-trained on the human genome. Except for the difference in the pre-training data, we kept all the pre-training setting the same as the DNABERT and trained a DNABERT model on the randomly generated nucleotide sequences. Through the contrastive comparison, we can investigate the first sub-task, as no biological information is contained in the totally random sequences for predicting context-dependency nucleotides. On the model level, we introduced randomness into the encoding modules of DNABERT, as shown in Figure 2. We re-initialized the learned weights in the encoding modules while only keeping learned k-mer embeddings of a pre-trained model. The “ablated” pre-trained model was further investigated in downstream tasks to compare with the other pre-trained models using human genome and totally random sequences. The model decomposition and dysfunctional analysis can be used for illustrating the learning the product of a pre-trained DNABERT model.

**Fig. 2:**
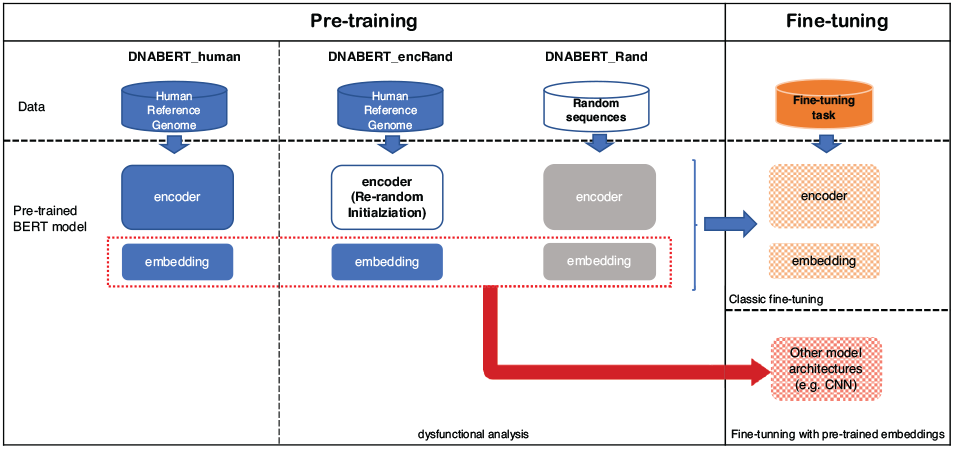
DNABERT model decomposition and evaluation strategy. The pre-trained model is decomposed into embedding and encoding modules. We use a dysfunctional approach to investigate the BERT model by incorporating randomness on both data and model levels. On the data level, we prepare totally randomly generated nucleotide sequences for around 3 billion lengths to compare with the DNABERT pre-trained on the human genome. On the model level, we introduce randomness into encoding layers through weights reinitialization. In the fine-tuning phage, besides the standard fine-tuning, we also evaluate only using different learned k-mer embeddings.

In addition, we also investigated using previously reported state-of-the-art but relatively “simpler” model structures (Deepromoter (Oubounyt *et al*., 2019) and CNN (Zeng *et al*., 2016)) for encoding layers with the pre-trained k-mer embeddings. Besides the two k-mer embeddings learned by DNABERT, we compared them with two other commonly used k-mer representations of dna2vec and one-hot embedding.

## 3 Results

### 3.1 Context-consistent k-mer representation is the major product learned by DNABERT in the embedding layer

As a k-mer can be literally decomposed into nucleotides, it is intuitive to evaluate the learned k-mer embeddings according to their nucleotide compositions. We visualized k-mer embeddings learned by DNABERT through tSNE analysis (Scikit-learn 1.1.1). For the pre-trained model on the human genome, we used the models provided by DNABERT (https://github.com/jerryji1993/DNABERT, abbreviated as DNABERT_human) directly. These models were trained on the human reference genome (GRCh38), and the weights of k-mer token embedding were extracted for tSNE analysis. For embeddings, we focus on k-mer token and ignore other embeddings, such as (token) type embedding. The type embedding is used for the specific NLP task of predicting the next sentence, which is not used in the pre-training task on nucleotide sequences. For the pre-trained model on random sequences (DANBERT_randData), we used DNABERT with default model parameters for pre-training. The k-mer representation learned from random sequences is extracted from the trained model. Besides the k-mer embeddings learned by DNABERT, we also visualized other commonly used k-mer embeddings of dna2vec (Ng, 2017) and one-hot embedding for comparison (No special token is included in the dna2vec and one-hot embedding).

We first investigated learned k-mer embeddings through tSNE visualization. Figure 3(a) and (b) demonstrate the 5-mer embeddings learned by the DNABERT model using different pre-training data. In the figures, we can observe the following: First, the special tokens are separated from other k-mer tokens in the space. Second, for k-mer tokens, k-mers with the same prefix or suffix nucleotide strings tend to cluster together. For example, CTCCN, NGCTT, and NGAAT (N is used as the abbreviation of A/T/G/C) in Figure 3(a) and (b). The local clustering of the same prefix and suffix strings results in a larger cluster group according to the center nucleotide (e.g., -GAT-). Therefore, we colored each k-mer according to the center position nucleotide (–[A/T/G/C]–) for easier viewing. In both Figure 3(a) and (b), there are four apparent clusters representing the nucleotide in the center position of a k-mer. **Although the randomly generated sequences do not contain any biological information, the context constraints of surrounding k-mers are still preserved in the random data**. Compared with the DNABERT trained on the human genome, k-mer embeddings of the DNABERT trained on randomly generated sequences are lined up more obviously. **Here, we refer to the adjacency of k-mer embeddings with a similar prefix or suffix as a context-consistent k-mer representation**. As a similar pattern can be observed in the k-mer representations of the pre-trained model usinga real genome and random nucleotide sequences, the context-consistent k-mer representation could be treated as one of the products that the DNABERT model learns in the embedding layer through pre-training. Such context-consistent k-mer representation is species-independent, which explains the generalization of the pre-training model across different species to some extent. For comparison, we also visualized the two commonly used k-mer embeddings of dna2vec (Ng, 2017) and one-hot encoding, shown in Figure 3(c) and (d). Although it is not as evident as in Figure 3(a) and (b), the closeness of the same prefix and suffix strings can also be observed in the dna2vec embedding. For example, NGAGC and GATGN are shown in Figure 3(c). No larger cluster according to the center position is not observed in the tSNE plot of dna2vec embedding. For the one-hot embedding, shown in Figure 3(d), k-mers are equidistant and evenly distributed.

**Fig. 3:**
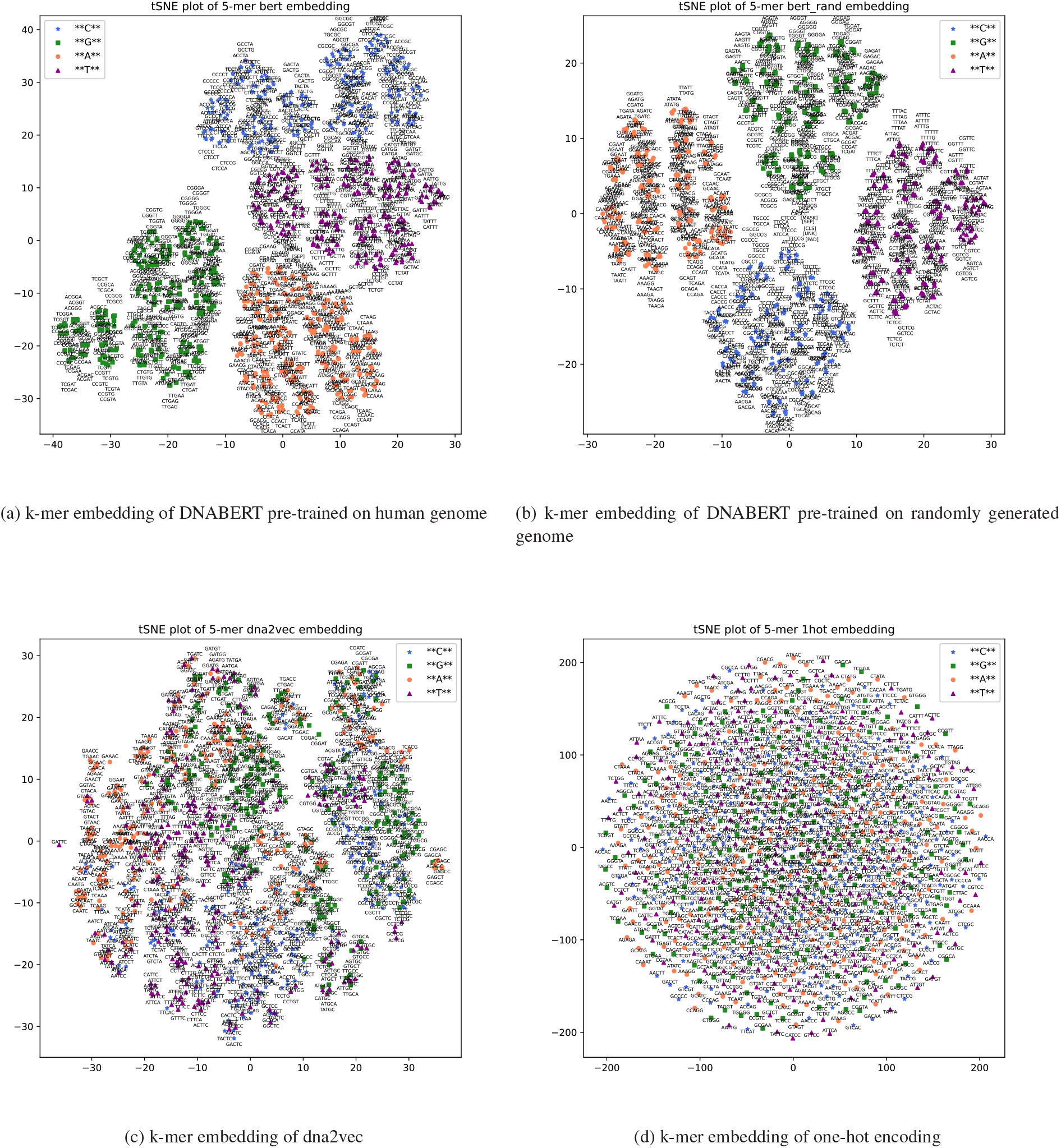
tSNE plot of 5-mer embeddings of DANBERT, DANBERT_dataRand, dna2vec and one-hot embedding. DNABERT_human is the DNABERT provided model pre-trained on human reference genome. DANBERT_dataRand is pre-trained on randomly generated sequences.

We then evaluated different DNABERT models and the use of different k-mer embeddings in the downstream tasks of sequence-based functional predictions, including TATA promoter prediction and transcript factor binding site (TFBS) prediction. We first evaluated the three pre-trained DNABERT models on the TATA promoter prediction task. Following the previous works (Oubounyt *et al*., 2019; Ji *et al*., 2021), the human and mouse positive promoter sequences were downloaded from the EPDnew database. These sequences are from the regions between -249 bp and 50 bp surrounding the transcription start site (TSS) position. In total, we have 29598 positive human samples (TATA:3065, nonTATA:26533) and 25110 positive mouse samples (TATA:3305, nonTATA:21805) with a sequence length of 300 bp. Negative sequences were generated accordingly to the positive samples. For TATA-containing samples, negative samples were selected from other genomic regions containing TATA-motif to avoid a more straightforward classification based on the presence of TATA-motif. For the non-TATA sequences, the positive samples were generated with random substitution of subsequences. We randomly split the dataset into the train, development, and test set (proportion 80/10/10).

The performance of different DNABERT models is shown in Figure 4 (or Supplementary material Table 1). Overall, on both datasets (human and mouse), DNABERT_human performs better than DNABERT_dataRand and DNABERT_encRand on evaluation metrics of AUROC, AURPRC, F1, and MCC. The performance gaps between DNABERT_human and DNABERT_dataRand, such as AUROC and AURPRC, are not very large (less than 0.0035). The performance gaps are relatively more significant on non-TATA data than on TATA data. The performance of DNABERT_dataRand and DNABERT_encRand are relatively close. Compared with DNABERT_human, DANBERT_encRand has the same pre-trained weights in the embedding layer, while pre-trained weights in the encoding layers are different. The performance gap between DNABERT_human and DNABERT_encRand demonstrates the role of encoding layers in the pre-training, which learns the combination of lower encoding layers or embeddings. The DNABERT_dataRand is trained on random data that contains no biological information in the sequences. The encoding layers do not learn any useful context constraint in the DNABERT_dataRand. Therefore, the learning product of the randomly re-initialized encoding layer in DNABERT_encRand can be approximately equivalent to the one in the encoding layers learned by the DNABERT_dataRand. The similar performance between DNABERT_encRand and DNABERT_dataRand indicates that the different part between learned k-mer embeddings using human and random data does not affect the fine-tuning performance much.

**Fig. 4:**
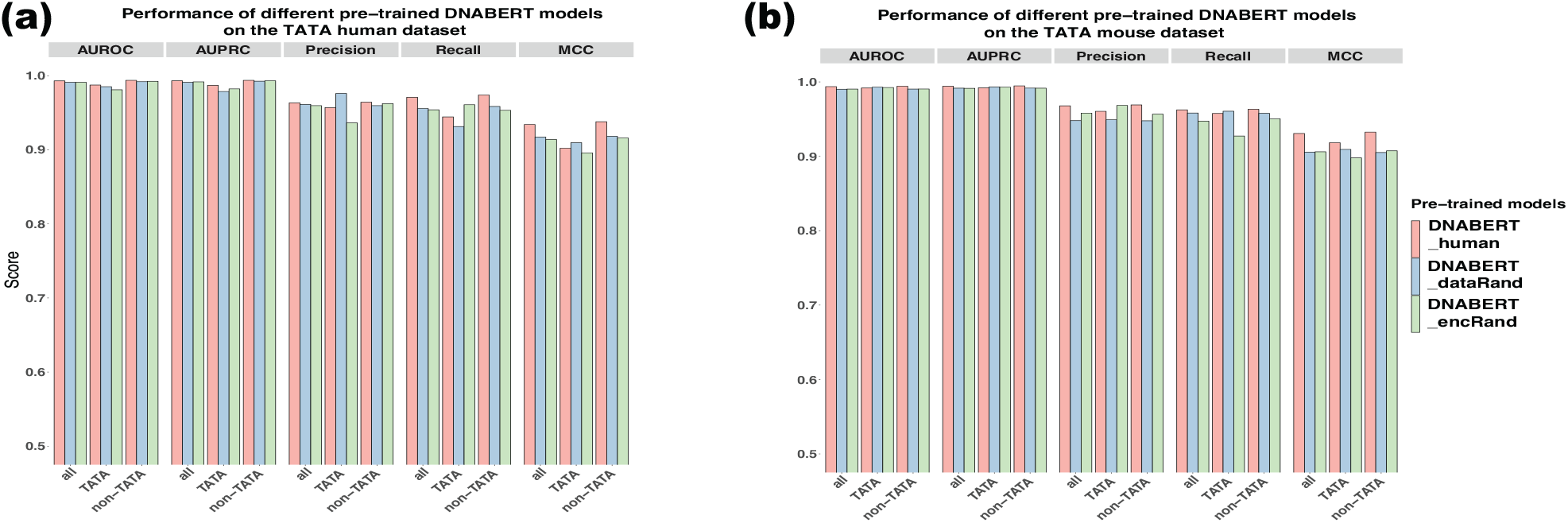
Performance of different pre-trained DNABERT models on the TATA human (a) and mouse dataset (b). DNABERT_human is the provided model pre-trained on human reference genome. DANBERT_dataRand is pre-trained on totally randomly generated sequences. DANBERT_encRand is based on DNABERT_human while the encoding layers are randomly re-initialized.

### 3.2 Evaluating different k-mer embeddings on TATA task

As the number of DNABERT’s model parameters is very large and the encoding layers take up 98% of the model parameters, we then investigated using simpler encoding layers in the downstream fine-tuning tasks. We evaluated four different k-mers embeddings and made use of Deepromoter (Oubounyt *et al*., 2019) as the basic framework for the TATA prediction task. Compared with DNABERT, Deepromoter has much fewer model parameters (shown in Figure 5(c) or in Supplementary material Table 2), while it is reported to achieve a good performance. We evaluated one-hot embedding (from 1-mer to 5-mer), dna2vec embedding (4-mer and 5-mer), and 5-mer embeddings of DNABERT using different pre-training datasets (human genome and random data). Unlike the default DNABERT fine-tuning, the extracted weights of the embedding layer learned from pre-training do not update anymore in the Deepromoter framework. We performed hyperparameters search on the development set of human data.

Shown in Figure 5(a) and (b), Deepromoter using embeddings learned by DNABERT have a better performance than it using the other two types of embeddings of one-hot and dna2vec. For the one-hot encoding, the number of k-mers increases exponentially as the k increases, which brings about the issue of sparse-high-dimensionality. As k increases, the one-hot encoding model reaches a performance peak when k equals three and starts to decline as k further increases. The best one-hot k-mer model performance is still lower than that achieved using k-mer embeddings of dna2vec and DNABERT. Dna2vec learned k-mer embeddings through a word embedding model word2vec trained on a two-layer neural network. In the provided dna2vec model, variant k-mer lengths are modeled, and different k-mer embeddings have the same dimension of 100. Although the k-mer dimension of dna2vec is smaller than that of DNABERT (768), the performances on AUROC, AUPRC and MCC are slightly worse than the model using k-mer embeddings of DNABERT. For the Deepromoter model only using k-mer embeddings learned by DNABERT, although the number of model parameters is much smaller than the DNABERT model, both performances (human 5-mer and random 5-mer) are close to the performance achieved by DNABERT_dataRand and DNABERT_encRand on the human dataset (slightly lower on the mouse dataset). When only using k-mer embeddings in the downstream TATA prediction task, the k-mer embedding of DNABERT trained on randomly generated data demonstrate its usability. The Deepromoter using DNABERT k-mer embedding trained on random data even achieves a better MCC result than the one trained on human data, which is more significant on the mouse dataset. Integrating with the tSNE plot in Figure 3, the context-consistent context k-mer embeddings from DNABERT can be treated as the major learning product in the embedding layer of the DNABERT model. The k-mer embedding of DNABERT itself can also be used independently as an alternative embedding choice to dna2vec and one-hot embedding for nucleotide sequence representation.

**Fig. 5:**
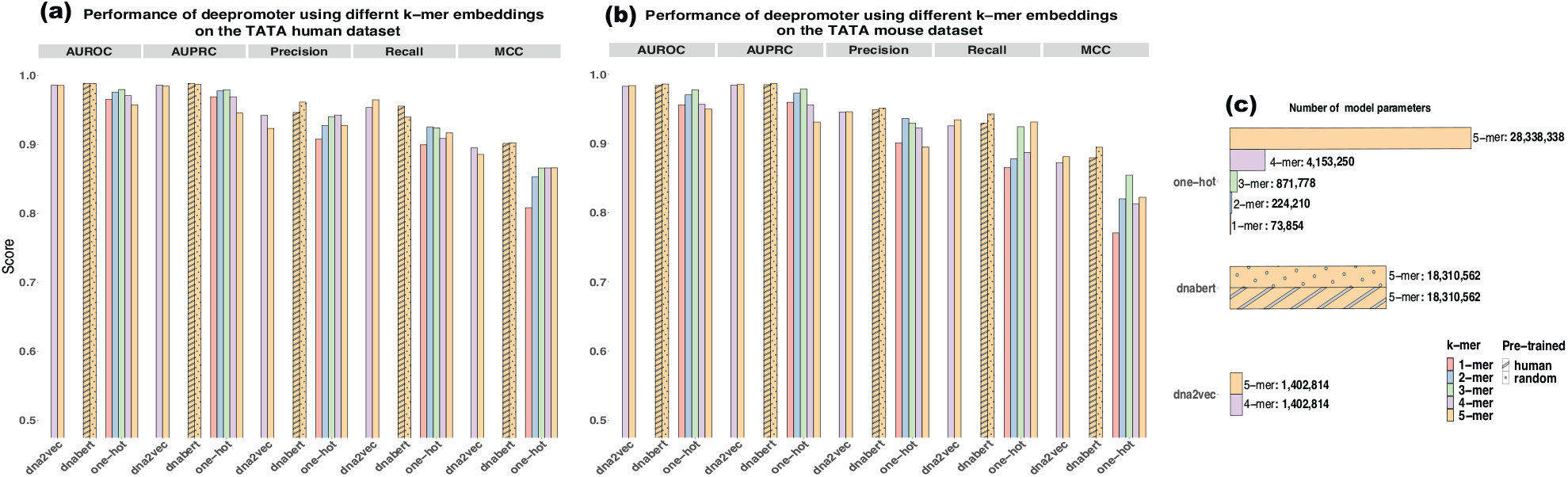
Performance of Deepromoter model using different k-mer embeddings on the TATA human (a) and mouse datasets (b). (c) Parameter numbers of the model using different k-mer embeddings.

### 3.3 Evaluation on TBFS tasks

We further evaluated the learned k-mer embeddings on TBFS tasks. The TFBS tasks include motif discovery, and motif occupancy identification (Zeng *et al*., 2016). Different from motif discovery, in which negative samples are shuffled positive sequences with the same dinucleotide frequency, the motif occupancy identification task uses negative samples that are also from regions centering at a motif instance with matched size, GC and motif strength. There are 690 motif discovery datasets and 422 motif occupancy datasets with input nucleotide sequences of 101 bp, which is less than the input sequence length of 300 bp in TATA promoter prediction task. For each dataset of ChIP-seq experiments, 80% of the data is used for training, and the rest 20% is used for testing. In each training dataset, 12.5% of the data is randomly selected and used as the development set. We used the classic CNN model (Zeng *et al*., 2016) as a simplified encoder for evaluating different k-mer embeddings. DNABERT models with fine-tuning are also evaluated for comparison. Basically, previously used hyperparameters of CNN (Zeng *et al*., 2016) and DNABERT were used as the default model parameters. Meanwhile, we randomly sampled 20 datasets to further tune model hyperparameters on the development set. On each training set, we trained 20 epochs and saved the model with the best MCC score on the development set. These saved models were used for the final predictions on the corresponding test set.

Figure 6 shows AUROC scores of the convolutional neural network (CNN) models using different k-mer embeddings and different DNABERT models with fine-tuning on 690 motif discovery datasets. We focused on the median AUROC score of the 690 datasets as an overall evaluation metric for different models. Other evaluation metrics, such as MCC, AUPRC, and F1, are shown in the Supplementary material. For one-hot embedding, we evaluated k-mer with lengths of 1, 4, and 5. The one-hot embeddings of 4-mer and 5-mer are used to compare with other learned k-mer embeddings with the vocabulary size in a similar scale range. One-hot embedding of 1-mer achieves a median AUROC score of 0.8551, which is consistent with the previously reported result (Zeng *et al*., 2016). When k increases to four and five, the dimension of the input vector increases to 256 and 1024, accordingly. The median AUROC score increases to 0.8929 and 0.8919 for 4-mer and 5-mer. The median AUROC scores are close to the one using dna2vec embedding (0.8918) with a vector dimension of 100. The median AUROC scores of the CNN model using DNABERT’s pre-trained embeddings are also very close, which are 0.8991 and 0.8965 for k-mer embedding pre-trained on human data and random sequence data. The performance of DNABERT fine-tuning models is shown in the last three columns in Figure 6. DNABERT achieves the best median AUROC score of 0.9. DNABERT_encRand and DNABERT_dataRand have similar performances of around 0.8652 and 0.8624, which are lower than CNN models using DNABERT’s k-mer embeddings. This performance gap illustrates the effect of the encoding module in the downstream fine-tuning. As the sizes of the 690 datasets are highly variant from 139 to 143425 (training data, median 21929.5), the complicated encoding module with large model parameters is hard to train with only using the fine-tuning dataset, especially on these datasets in small scales. Therefore, sufficient pre-training becomes a pre-requisition when using a complicated encoding module. On the other hand, we observe that the performance gap between CNN with embeddings and the DNABERT fine-tuning model is not very significant.

**Fig. 6:**
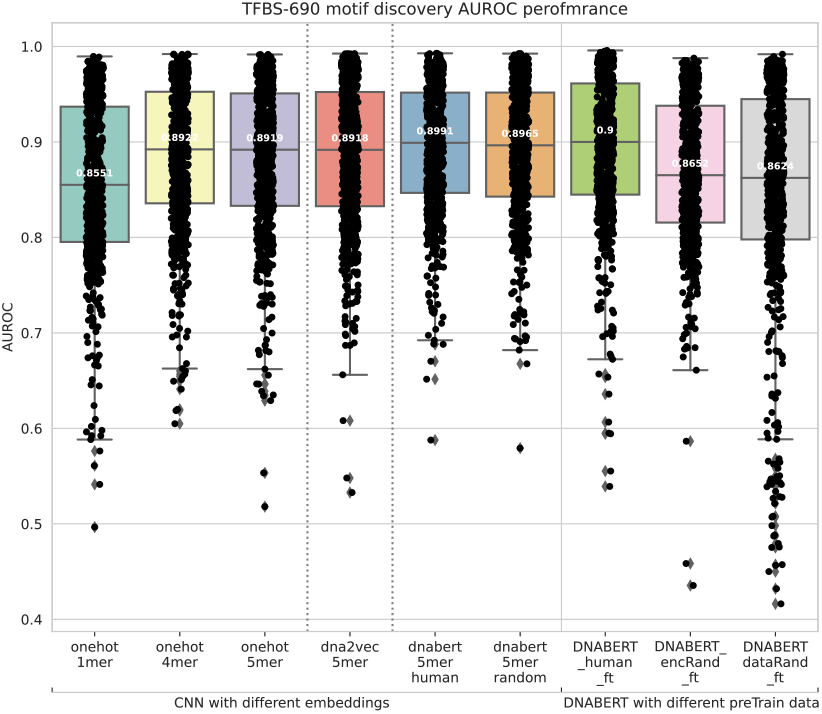
Boxplot of AUROC performance on 690 TFBS motif discovery datasets.

We also performed a similar evaluation on the 422 TFBS motif occupancy identification datasets (Figure 7). As the motif occupancy identification is more difficult than the motif discovery task (Zeng *et al*., 2016), the median AUROC score generally has a lower value than the one in the motif discovery task, ranging from 0.80 to 0.8465. For the models using the same type of k-mer embeddings, a similar performance pattern can be observed as in the motif discovery task. One different but interesting result is that the CNN using 5-mer one-hot embedding achieves the best AUROC median score of 0.8465, which is even slightly better than the DNABERT fine-tuning model of 0.8398. Although confounding factors (e.g., different model parameters) exist, this indicates that a smaller neural network structure could be more proper for applications on a relatively small-scale dataset.

**Fig. 7:**
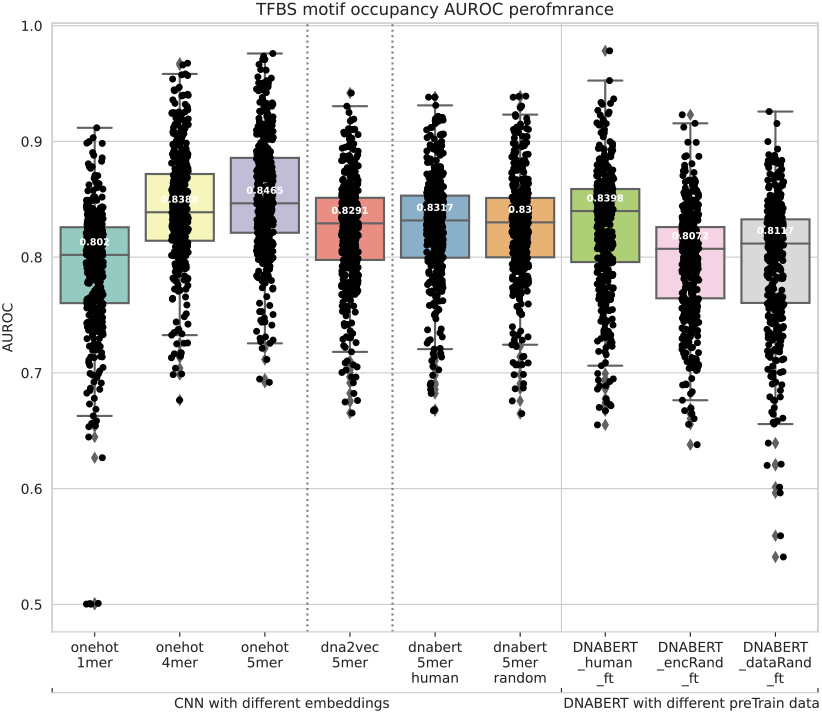
Boxplot of AUROC Performances on 422 TFBS motif occupancy identification datasets.

### 3.4 Pre-training can be accelerated through model initialization with the model weights trained on random data

Pre-training BERT models are known to be computationally intensive and time-consuming. Although a pre-trained model can be applied cross-species, this assumes species that are not far distance away in the phylogenetic tree. For example, for a downstream application related to bacteria genomes, it could be better to use a pre-trained model based on bacteria data than directly using a pre-trained model based on human data, considering the role of the encoding module. For the need to pre-train a new model, the random data pre-trained model can be used to initialize model weights for accelerating the pre-training process, considering the context-consistent k-mer embedding has already been learned.

Figure 8 demonstrates the exponential loss on the development set of the pre-training with and without model initialization on the human reference genome. Pre-training with the model initialization takes around 1500 iterations to get into the stage of convergence, while pre-training without the model initialization starts to get into coverage after 5500 iterations. As the pre-training on random sequence data is non-specie-specific, no additional bias is introduced in the new pre-trained model.

**Fig. 8:**
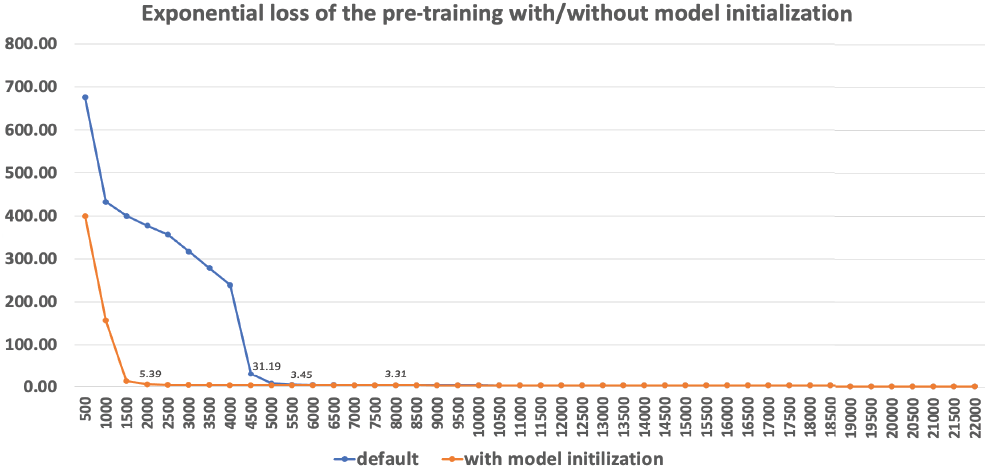
Exponential loss curve on the development set of the pre-training with and without model initialization. The development set is randomly selected 20% of the total training data of human reference genome (hg38). Performing model initialization with the weights of the model trained on random data can reduce the number of iterations for convergence.

## 4 Discussion

For protein sequences, previous work (Yang *et al*., 2018) reports that randomizing the UniPort sequences improves downstream performance for some tasks, such as cytochrome P450 thermostability. They suspected this relates to the regularizing effect on the embedding model. The randomization prevents the embedding model to a set of protein sequences in the pre-train dataset. Here, we analyzed from a different perspective that aims to illustrate the learning product of the DNABERT model in the embedding layer. The other difference is that the k-mers here are overlapped, while the amino acid sequences can be treated as non-overlapped k-mers. We demonstrated that the consistent-context k-mer embeddings are the primary product learned by the BERT model in the embedding layer. Through two downstream tasks, we showed that embeddings learned from random nucleotide sequence data can also be helpful in downstream tasks in the pre-training and fine-tuning paradigm. The embedding learned from random data can perform similarly to the embeddings learned on actual genome data. This also explains the generalization ability in the embedding layer from a different perspective. Moreover, the k-mer embedding learned on the random data can provide an unbiased choice of k-mer embedding for various downstream tasks.

One commonly used approach to illustrate the effect of pre-training is training from scratch. In this approach, the pre-training step is skipped, and the model is directly trained with fine-tuning. Such scratch-training usually brings about poor performance (Ji *et al*., 2021), as the model with a large number of parameters is difficult to be sufficiently trained only using a small amount of fine-tuning dataset. In this work, we introduced two dysfunctional approaches to illustrate the effect of pre-training. Besides introducing randomness into the encoding layers, we also conducted pre-training on totally random sequences. Compared with the scratch-training, using the pre-trained model on random sequences achieves relatively close results in TATA and TBFS tasks. This supports the observation that context-consistent embedding is the primary learning product in the embedding layer. Meanwhile, the pre-trained model on totally random data can provide a pivot to investigate the biologically insignificant perturbations of the input nucleotide sequences.

## 5 Conclusion

In this work, focusing on nucleotide sequences, we illustrated the learning product of a typical BERT model learned through pre-training. We used a dysfunctional approach to investigate different modules by incorporating randomness on both data and model levels. We demonstrated that context-consistent k-mer representation is the primary result of a typical BERT model learns in the embedding layer and the effect of encoding layers in downstream tasks. We compared the learned k-mer embeddings with other commonly used k-mer representations in downstream tasks of sequence-based functional prediction. In addition, we proposed a simple but effective approach to accelerate the pre-training process.

## Footnotes

1 In the following part, BERT and DNABERT are interchangeable, referring to the BERT model for nucleotide sequences.

