## Supplementary material for "Dysfunctional analysis of the pre-training model on nucleotide sequences and the evaluation of different k-mer embeddings"

### Supplementary materials

#### 1.Data

##### 1.1 Random sequence generation and pre-training data preparation

We repeatedly used the python (v3.8.10) function random.choice to generate random sequences. The random seed used in our experiment is 123. In total, we generated a random sequence of 3 billion nucleotides. Then, we used process_pretrain_data.py provided by DNABERT to prepare the pre-training data on randomly generated sequences.

#### 2. Models

##### 2.1 DNABERT fine-tunning models

We used the same model as used in the paper (Ji et al. Bioinformatics 2021).
https://drive.google.com/file/d/1KMqgXYCzrrYD1qxdyNWnmUYPtrhQqRBM/view
For fine-tuning, DNABERT uses the default learning rate of 1e-4. DNABERT_dataRand and DNABERT_encRand use a learning rate of 1e-5 to avoid training collapse.

##### 2.2 Deepromoter with different k-mers

###### 2.2.1 Hyperparameters

```
KMER=5
SPIECE="mouse/human"
MODEL="deepPromoterNet"
EMBEDDING="onehot/dna2vec/dnabert"
KERNEL="5,5,5"
LR=1e-4
EPOCH=20
DROPOUT=0.1
BS=64
```

###### 2.2.2 Model structure

```
deepPromoterNet(
  (cnn_diff_kernel): ModuleList(
    (0): Sequential(
      (0): Conv1d(16, 16, kernel_size=(5,), stride=(1,), padding=same)
      (1): ReLU(inplace=True)
      (2): MaxPool1d(kernel_size=6, stride=6, padding=0, dilation=1, ceil_mode=False)
      (3): Dropout(p=0.1, inplace=False)
```

```
    )
    (1): Sequential(
      (0): Conv1d(16, 16, kernel_size=(5,), stride=(1,), padding=same)
      (1): ReLU(inplace=True)
      (2): MaxPool1d(kernel_size=6, stride=6, padding=0, dilation=1, ceil_mode=False)
      (3): Dropout(p=0.1, inplace=False)
    )
    (2): Sequential(
      (0): Conv1d(16, 16, kernel_size=(5,), stride=(1,), padding=same)
      (1): ReLU(inplace=True)
      (2): MaxPool1d(kernel_size=6, stride=6, padding=0, dilation=1, ceil_mode=False)
      (3): Dropout(p=0.1, inplace=False)
    )
  )
  (biLSTM): LSTM(49, 32, batch_first=True, bidirectional=True)
  (linear): Linear(in_features=64, out_features=32, bias=True)
  (flatten): Flatten(start_dim=1, end_dim=-1)
  (fc): Sequential(
    (0): Linear(in_features=1536, out_features=128, bias=True)
    (1): ReLU()
    (2): Dropout(p=0.1, inplace=False)
    (3): Linear(in_features=128, out_features=2, bias=True)
    (4): Softmax(dim=1)
  )
)
 |- Total [deepPromoterNet] Parameters 224,210
```

**2.3 CNN (Zeng et al.,) with different k-mers**

**2.3.1 Hyperparameters**

```
MODEL="zeng_CNN"
KERNEL="24
EMBEDDING="onehot/dna2vec/dnabert"
LR=0.001
EPOCH=10
BS=64
dropout=0.1
```

**2.3.2 Model structure:**

```
zeng_CNN(
  (cnn_1m): Sequential(
```

```
    (0): Conv1d(voc_size, 128, kernel_size=(24,), stride=(1,), padding=same)
    (1): ReLU(inplace=True)
  )
  (fc): Sequential(
    (0): Linear(in_features=128, out_features=32, bias=True)
    (1): ReLU()
    (2): Dropout(p=0.1, inplace=False)
    (3): Linear(in_features=32, out_features=2, bias=True)
    (4): Softmax(dim=1)
  )
)
 |- Total [zeng_CNN] Parameters 3,150,050 (5-mer)
```

#### 2.4 Machine training GPU setting

DNABERT Nvidia V100 x 2
Deepromoter + different embeddings  Nvidia A6000 x 1
CNN + different embeddings  Nvidia A6000 x 1

For reproducibility, we set the random seed to be 123.

### 3. Supplemental Figures and Tables

#### 3.1 Supplemental Figures

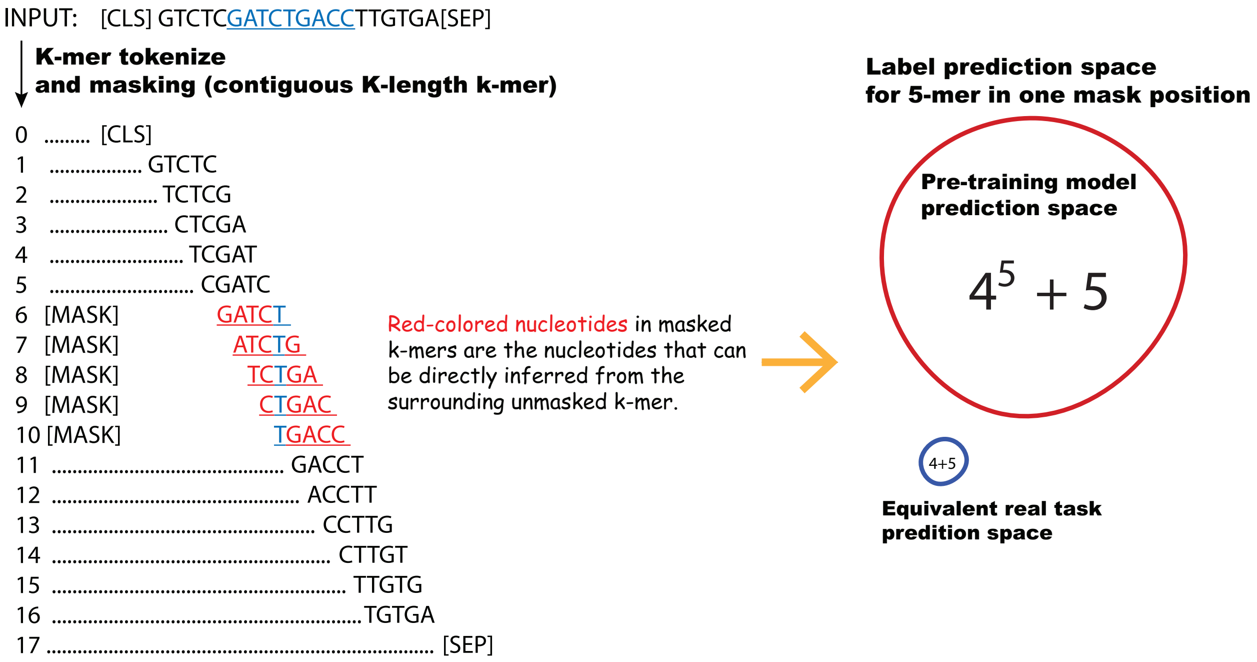


**Figure S1. The k-mer sequence masking strategy used in DNABERT. To avoid a masked token from being trivially inferred through the immediately surrounding k-mers, DNABERT masks k contiguous k-mers. Compared with the non-inferable nucleotides predicting sub-task, the original prediction space is huge.**

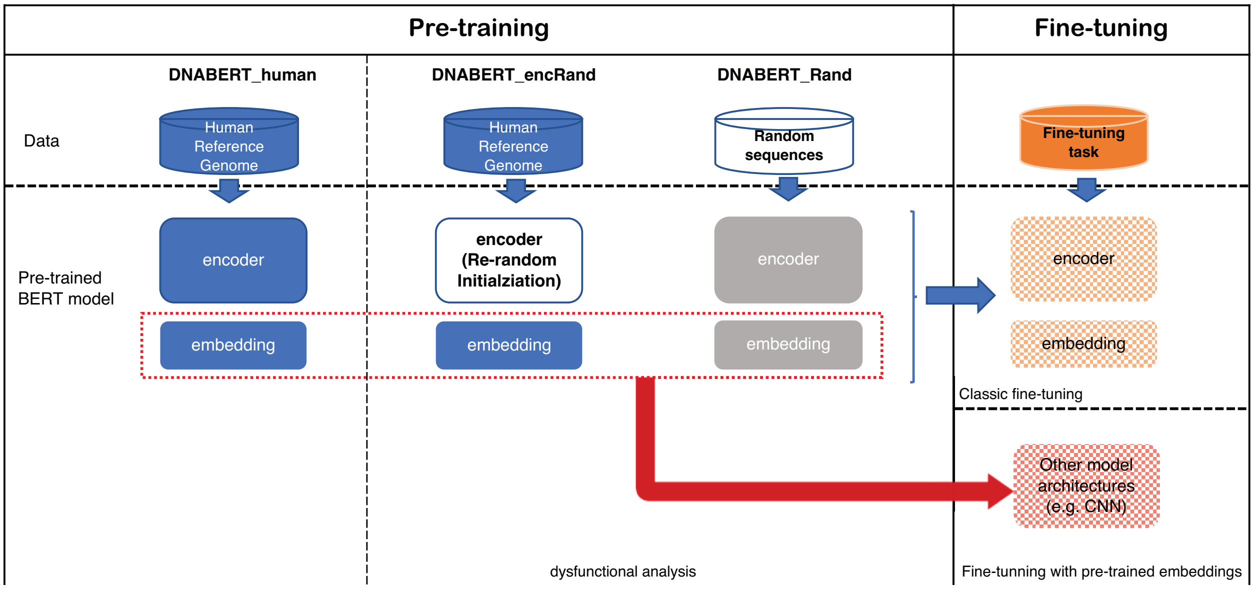


**Figure S2. DNABERT model decomposition and evaluation strategy. The pre-trained model is decomposed into embedding and encoding modules. We use a dysfunctional approach to investigate the BERT model by incorporating randomness on both data and model levels. On the data level, we prepare totally randomly generated nucleotide sequences for around 3 billion lengths to compare with the DNABERT pre-trained on the human genome. On the model level, we introduce randomness into encoding layers through weights reinitialization. In the fine-tuning phage, besides the standard fine-tuning, we also evaluate only using different learned k-mer embeddings.**

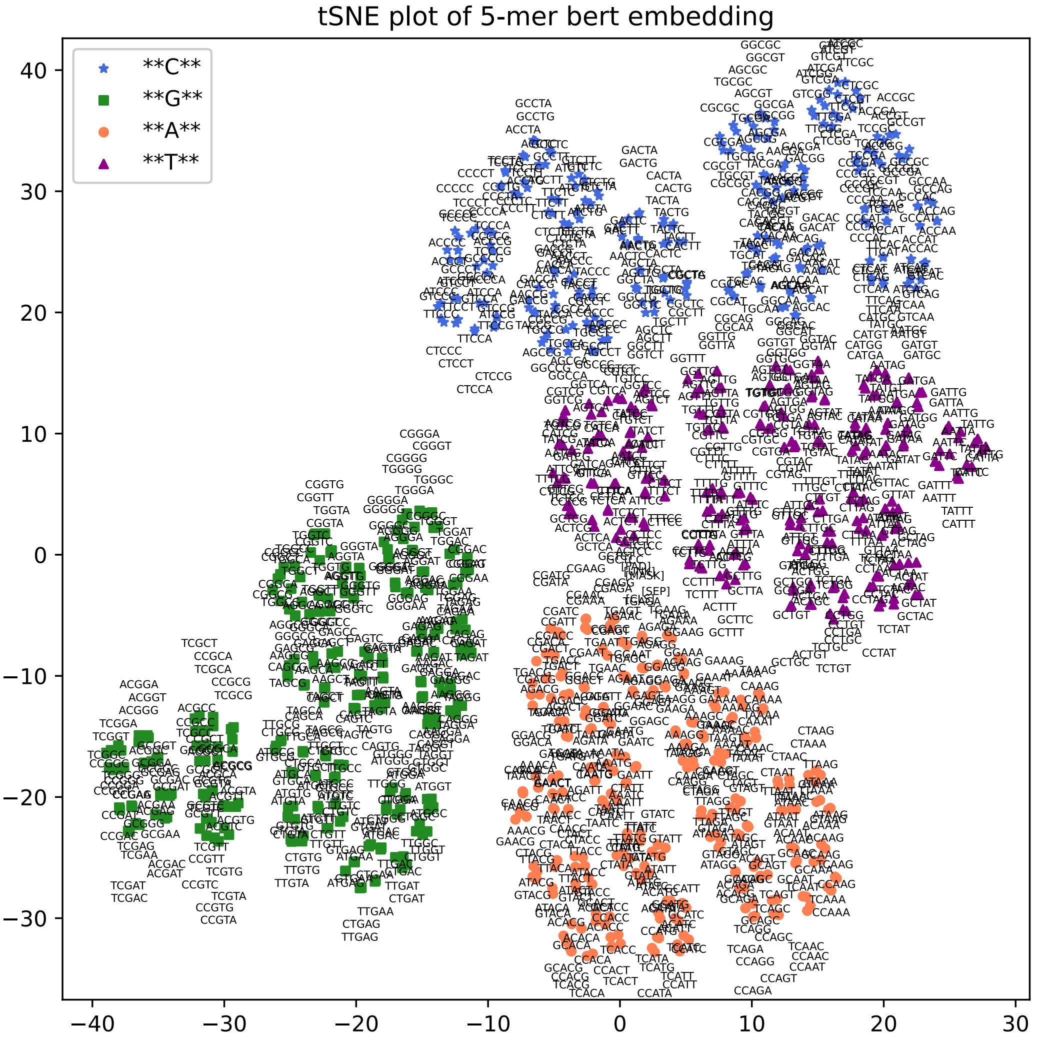


**Figure S3. tSNE plot of the 5-mer embedding of DNABERT (provided by Ji et al., 2021) pre-trained on human genome data.**

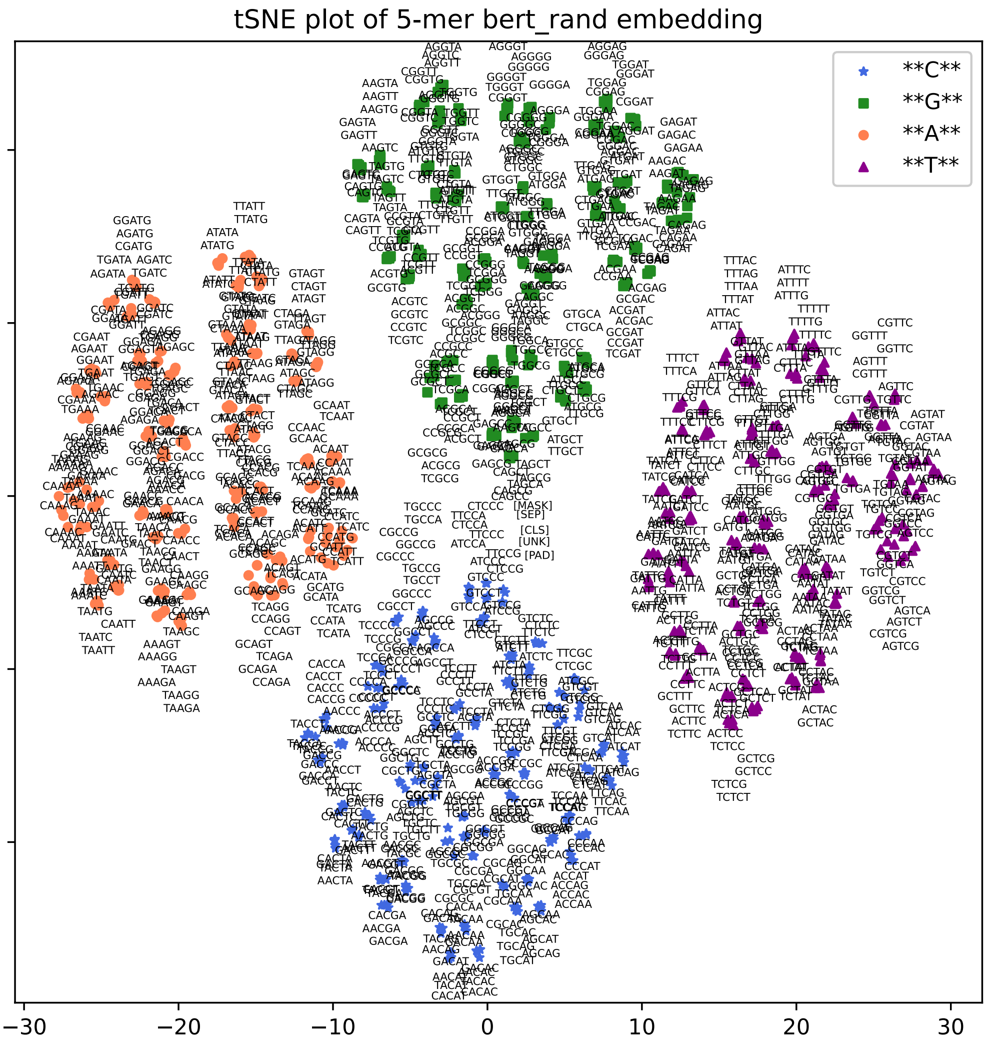


**Figure S4. tSNE plot of the 5-mer embedding of DNABERT pre-trained on totally random nucleotide sequences around 3 billion nucleotides.**

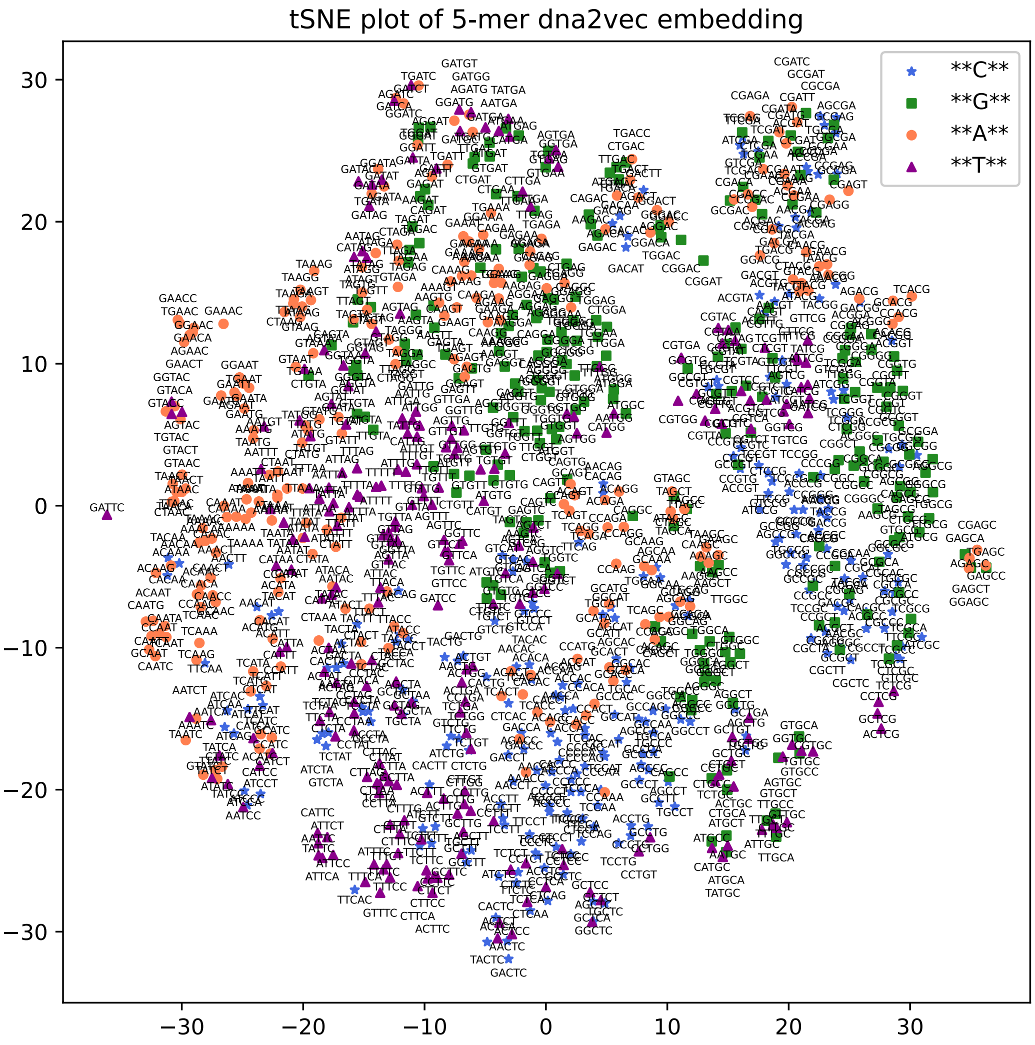


**Figure S5. tSNE plot of the 5-mer embedding of dna2vec (provided by Ng. et al., 2017).**

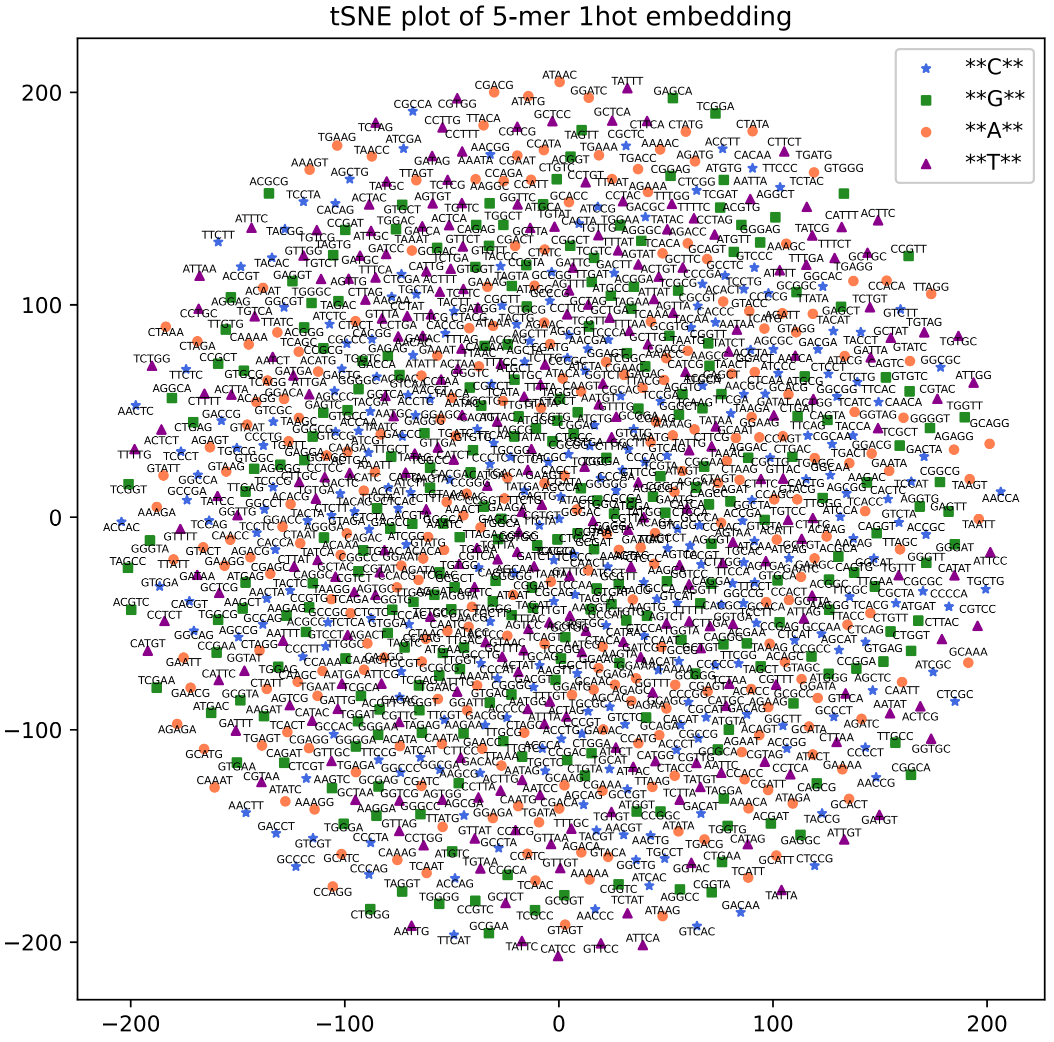


**Figure S6. tSNE plot of the 5-mer embedding of one-hot encoding.**

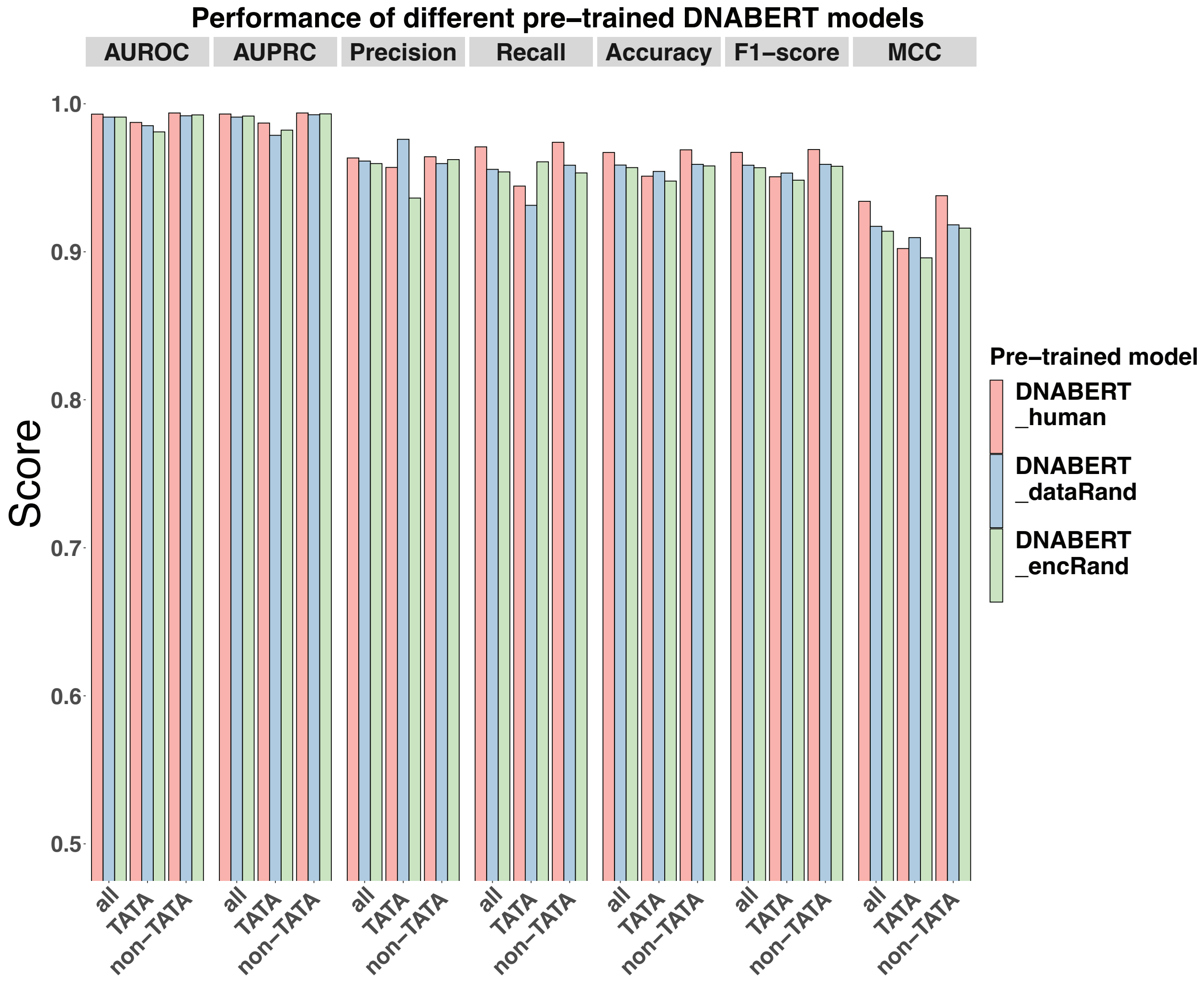


**Figure S7. Performance of different pre-trained DNABERT models on the TATA human dataset.**

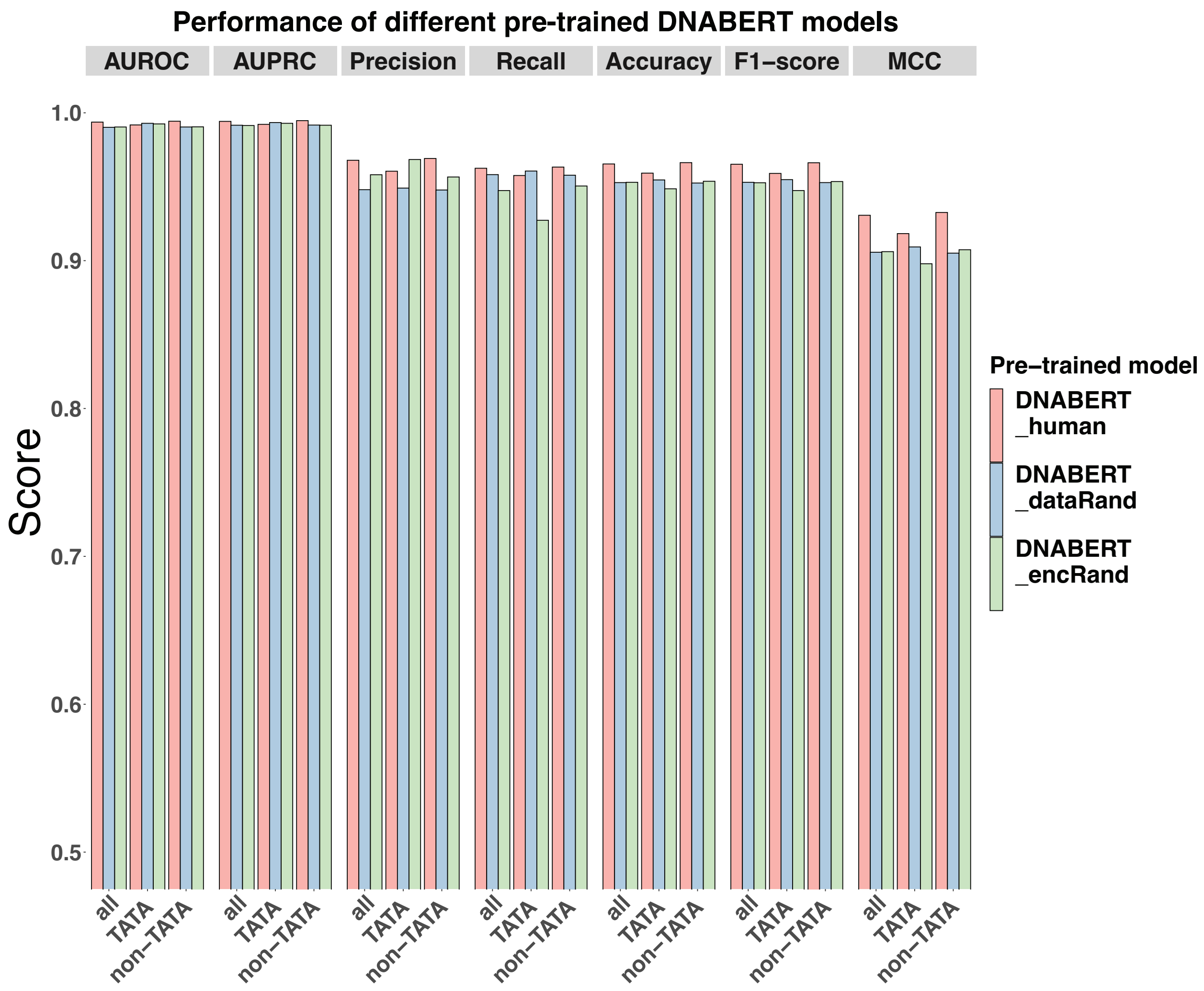


**Figure S8. Performance of different pre-trained DNABERT models on the TATA mouse dataset.**

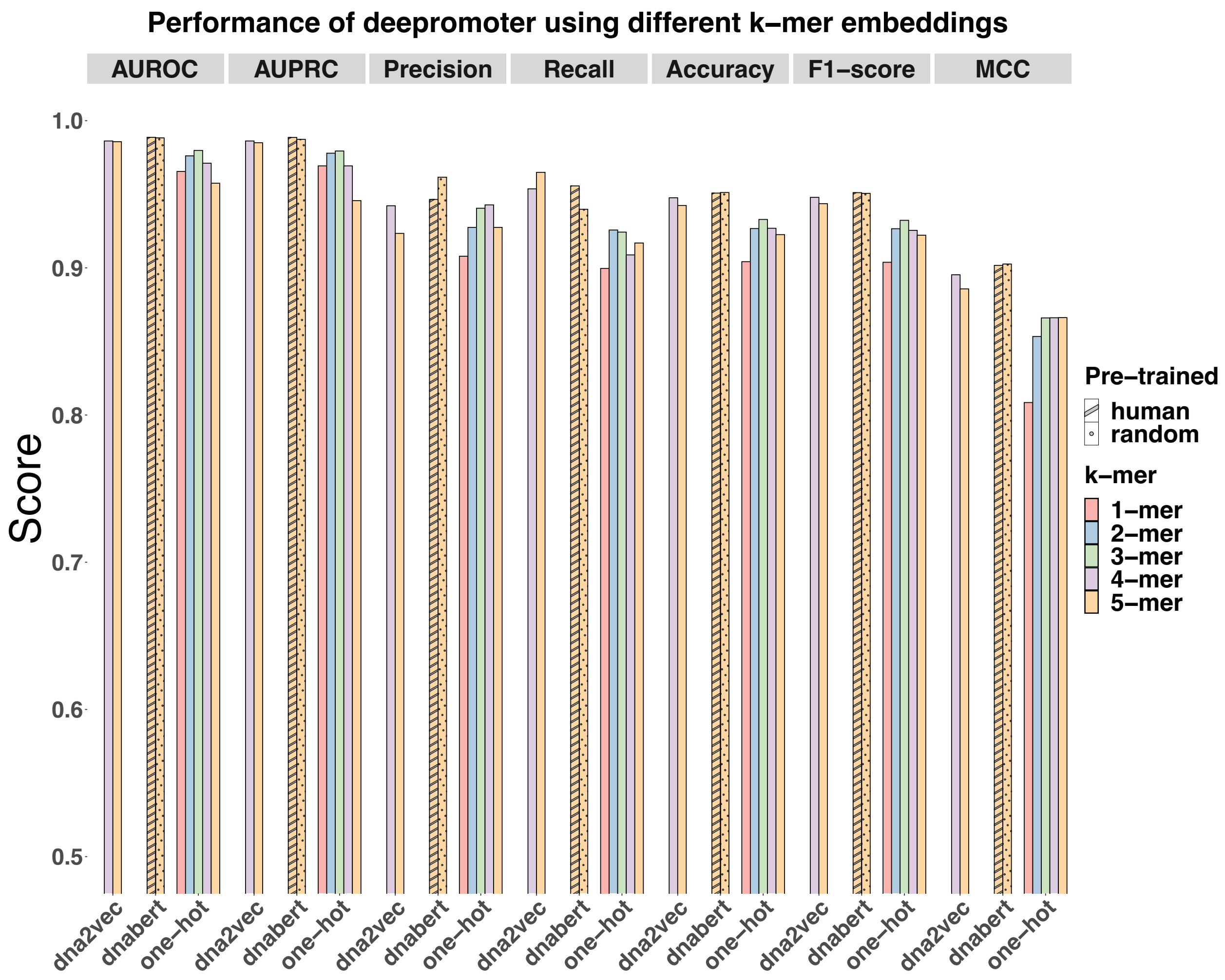


**Figure S9. Performance of Deepromoter model using different k-mer embeddings on the TATA human dataset.**

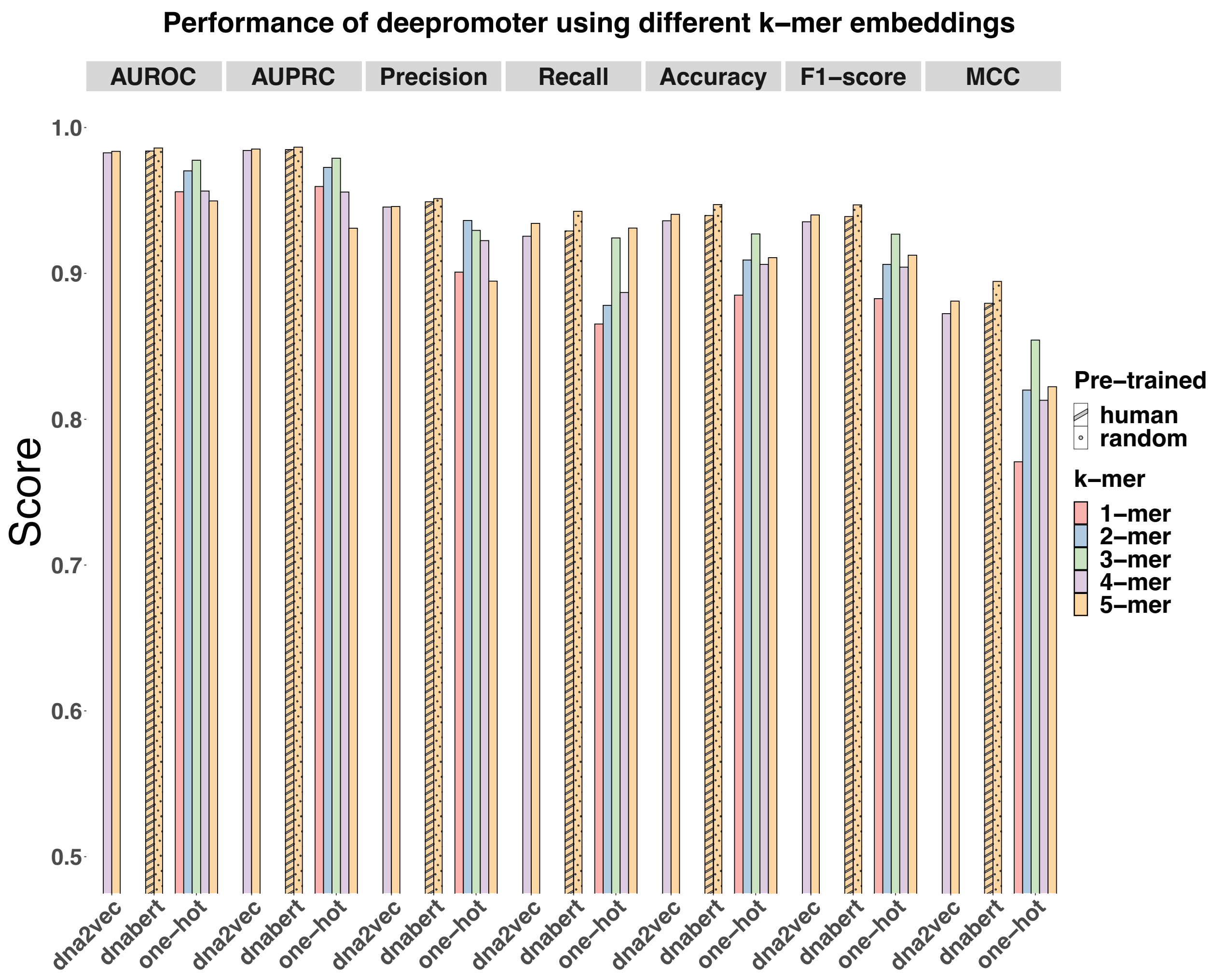


**Figure S10. Performance of Deepromoter model using different k-mer embeddings on the TATA mouse dataset.**

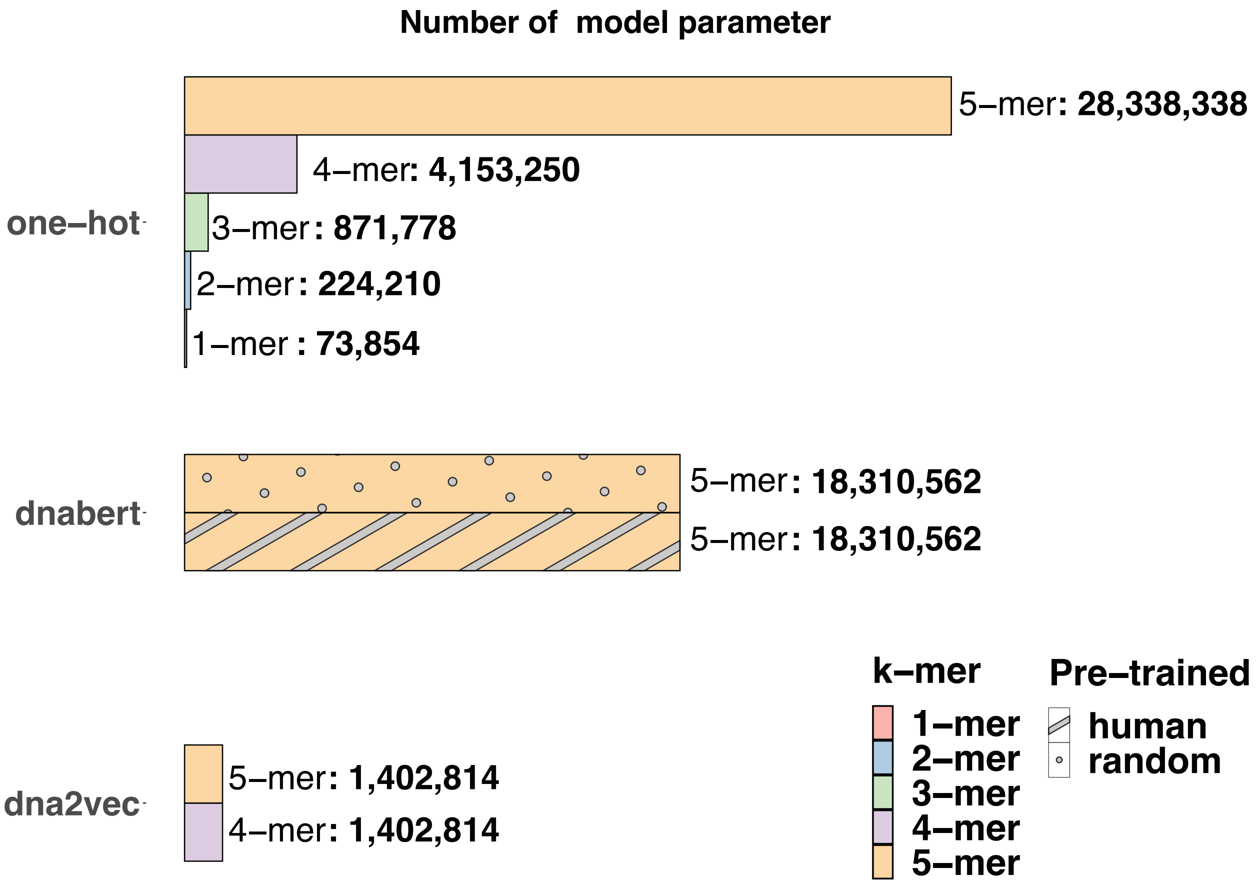


**Figure S10. Number of model parameters of Deepromoter using different k-mer embeddings.**

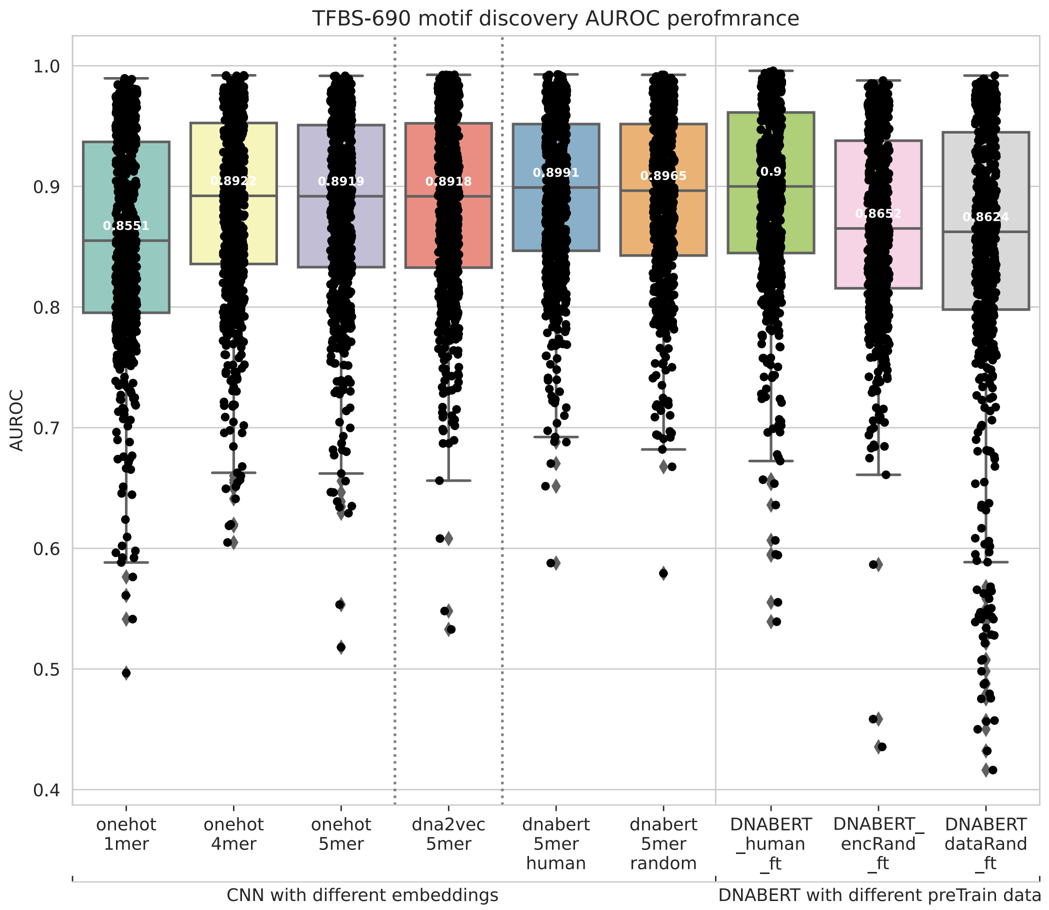


**Figure S11. Boxplot of AUROC performance on 690 TFBS motif discovery datasets.**

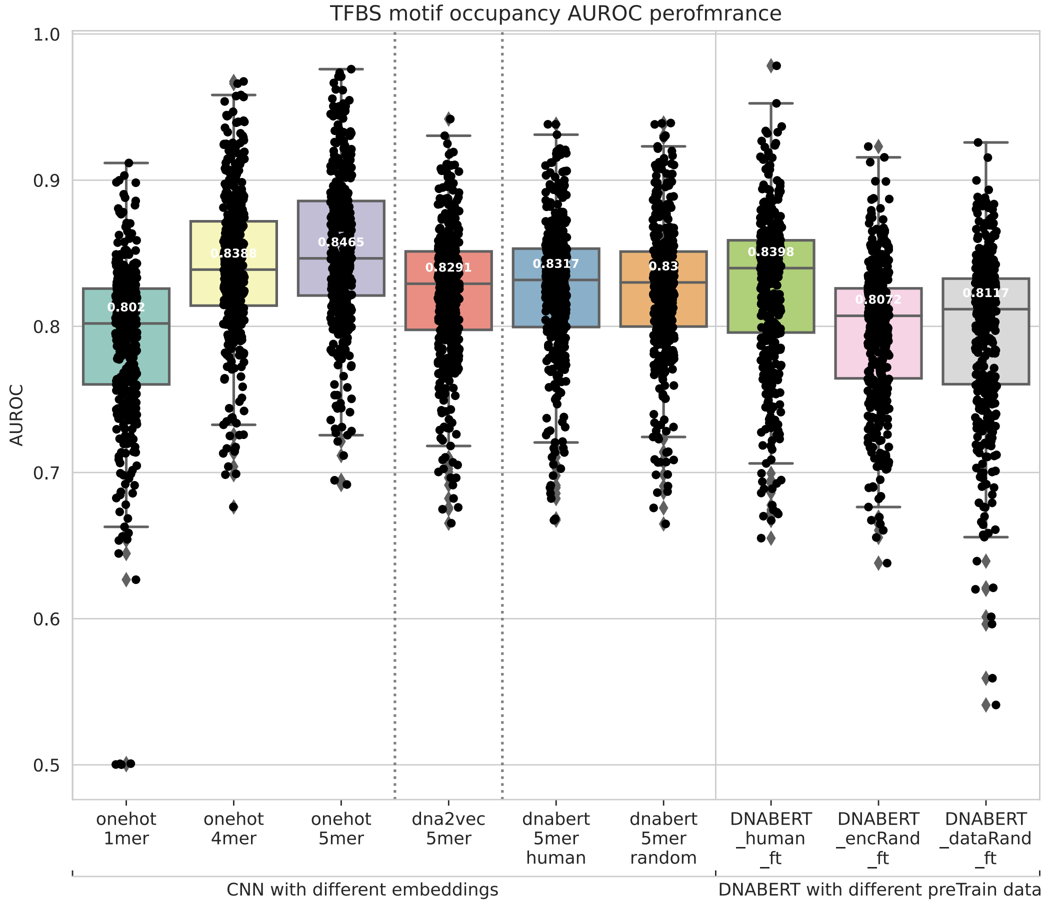


**Figure S12. Boxplot of AUROC Performances on 422 TFBS motif occupancy identification datasets.**

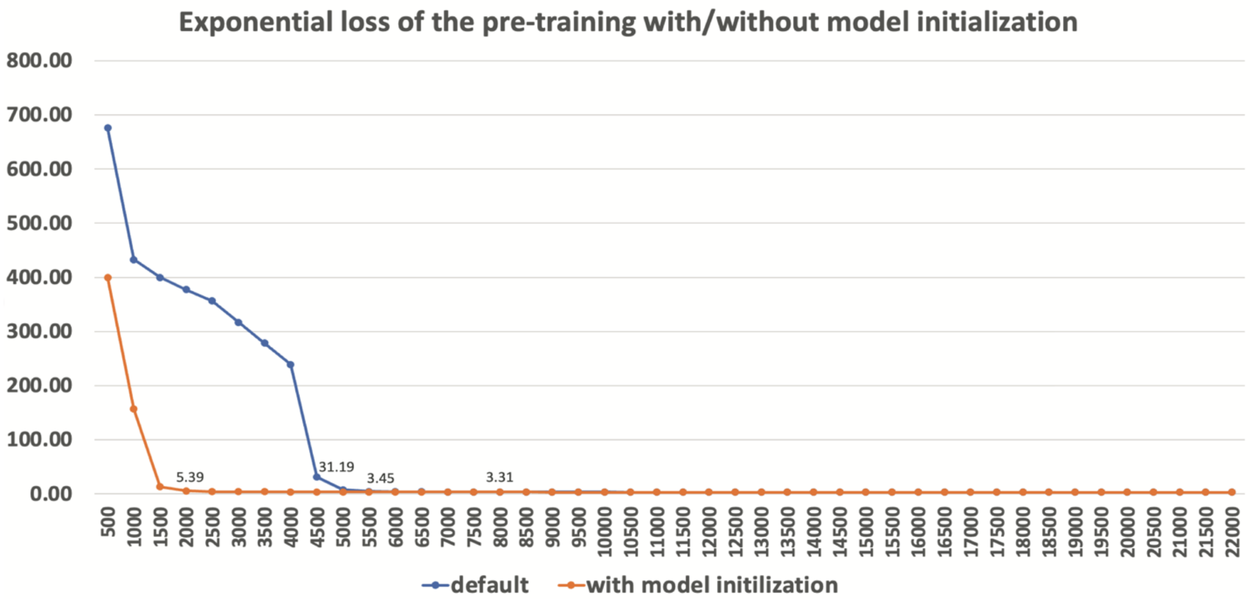


**Figure S13. Exponential loss curve on the development set of the pre-training with and without model initialization. The development set is randomly selected 20% of the total training data of human reference genome (hg38).Performing model initialization with the weights of the model trained on random data can reduce the number of iterations for convergence.**

##### 3.2 Tables

**Human dataset**

| Pre-trained model | Test data category | AUROC | AUPRC | Precision | Recall | Accuracy | F1-score | MCC |
|---|---|---|---|---|---|---|---|---|
| DNABERT_human | all | **0.9930** | 0.9931 | 0.9634 | 0.9709 | 0.9671 | 0.9672 | **0.9341** |
| | TATA | **0.9874** | 0.9870 | 0.9570 | 0.9444 | 0.9511 | 0.9507 | 0.9022 |
| | non-TATA | **0.9938** | 0.9938 | 0.9642 | 0.9740 | 0.9689 | 0.9691 | **0.9379** |
| DNABERT_dataRand | all | 0.9910 | 0.9910 | 0.9613 | 0.9557 | 0.9586 | 0.9585 | 0.9172 |
| | TATA | 0.9852 | 0.9787 | 0.9760 | 0.9314 | 0.9543 | 0.9532 | **0.9096** |
| | non-TATA | 0.9919 | 0.9926 | 0.9596 | 0.9585 | 0.9591 | 0.9591 | 0.9182 |
| DNABERT_encRand | all | 0.9910 | 0.9917 | 0.9596 | 0.9540 | 0.9569 | 0.9568 | 0.9139 |
| | TATA | 0.9810 | 0.9822 | 0.9363 | 0.9608 | 0.9478 | 0.9484 | 0.8959 |
| | non-TATA | 0.9925 | 0.9932 | 0.9623 | 0.9533 | 0.9580 | 0.9578 | 0.9160 |

**Mouse dataset**

| Pre-trained model | Test data | AUROC | AUPRC | Precision | Recall | Accuracy | F1-score | MCC |
|---|---|---|---|---|---|---|---|---|
| DNABERT_human | all | **0.9937** | 0.9942 | 0.9679 | 0.9625 | 0.9654 | 0.9652 | **0.9307** |
| | TATA | 0.9918 | 0.9922 | 0.9605 | 0.9576 | 0.9592 | 0.9590 | **0.9183** |
| | non-TATA | **0.9943** | 0.9947 | 0.9691 | 0.9633 | 0.9663 | 0.9662 | **0.9326** |
| DNABERT_dataRand | all | 0.9902 | 0.9916 | 0.9480 | 0.9582 | 0.9528 | 0.9530 | 0.9057 |
| | TATA | **0.9929** | 0.9934 | 0.9491 | 0.9606 | 0.9546 | 0.9548 | 0.9093 |
| | non-TATA | 0.9904 | 0.9917 | 0.9478 | 0.9578 | 0.9525 | 0.9528 | 0.9051 |
| DNABERT_encRand | all | 0.9904 | 0.9914 | 0.9581 | 0.9474 | 0.9530 | 0.9527 | 0.9061 |
| | TATA | 0.9925 | 0.9929 | 0.9684 | 0.9273 | 0.9486 | 0.9474 | 0.8979 |
| | non-TATA | 0.9905 | 0.9916 | 0.9566 | 0.9505 | 0.9537 | 0.9535 | 0.9074 |

**Table 1.** Fine-tuning performance of different pre-trained DNABERT models on human and mouse dataset. DNABERT_human is the provided model pre-trained on human reference genome. DANBERT_dataRand is pre-trained on totally randomly generated sequences. DANBERT_encRand is based on DNABERT_human while the encoding layers are randomly re-initialized. For different test data, the best AUROC and MCC scores are shown in bold font.

**Human dataset**

| Embedding | k-mer | Voc size | Model parameter | AUROC | AUPRC | Precision | Recall | Accuracy | F1-score | MCC |
|---|---|---|---|---|---|---|---|---|---|---|
| one-hot | 1-mer | 4 | 73854 | 0.9655 | 0.9693 | 0.9079 | 0.8996 | 0.9042 | 0.9038 | 0.8085 |
| | 2-mer | 16 | 224210 | 0.9761 | 0.9779 | 0.9275 | 0.9257 | 0.9267 | 0.9266 | 0.8534 |
| | 3-mer | 64 | 871778 | 0.9798 | 0.9794 | 0.9405 | 0.9243 | 0.9329 | 0.9323 | 0.8660 |
| | 4-mer | 256 | 4153250 | 0.9711 | 0.9693 | 0.9428 | 0.9088 | 0.9269 | 0.9255 | 0.8543 |
| | 5-mer | 1024 | 28338338 | 0.9575 | 0.9457 | 0.9275 | 0.9169 | 0.9226 | 0.9222 | 0.8453 |
| dna2vec | 4-mer | 100 | 1402814 | 0.9862 | 0.9862 | 0.9422 | 0.9537 | 0.9476 | 0.9479 | 0.8953 |
| | 5-mer | 100 | 1402814 | 0.9857 | 0.9850 | 0.9234 | 0.9649 | 0.9424 | 0.9436 | 0.8857 |
| dnabert | human 5-mer | 768 | 18310562 | **0.9887** | 0.9886 | 0.9465 | 0.9557 | 0.9508 | 0.9511 | 0.9017 |
| | random 5-mer | 768 | 18310562 | 0.9883 | 0.9873 | 0.9616 | 0.9398 | 0.9512 | 0.9506 | **0.9026** |

**Mouse dataset**

| Embedding | k-mer | Voc size | Model parameter | AUROC | AUPRC | Precision | Recall | Accuracy | F1-score | MCC |
|---|---|---|---|---|---|---|---|---|---|---|
| one-hot | 1-mer | 4 | 73854 | 0.9560 | 0.9596 | 0.9009 | 0.8653 | 0.8851 | 0.8827 | 0.7708 |
| | 2-mer | 16 | 224210 | 0.9703 | 0.9727 | 0.9363 | 0.8781 | 0.9092 | 0.9062 | 0.8200 |
| | 3-mer | 64 | 871778 | 0.9776 | 0.9790 | 0.9295 | 0.9243 | 0.9271 | 0.9269 | 0.8543 |
| | 4-mer | 256 | 4153250 | 0.9565 | 0.9558 | 0.9225 | 0.8869 | 0.9062 | 0.9043 | 0.8130 |
| | 5-mer | 1024 | 28338338 | 0.9497 | 0.9310 | 0.8947 | 0.9311 | 0.9108 | 0.9125 | 0.8223 |
| dna2vec | 4-mer | 100 | 1402814 | 0.9827 | 0.9843 | 0.9455 | 0.9255 | 0.9361 | 0.9354 | 0.8724 |
| | 5-mer | 100 | 1402814 | 0.9837 | 0.9853 | 0.9459 | 0.9343 | 0.9405 | 0.9401 | 0.8810 |
| dnabert | human 5-mer | 768 | 18310562 | 0.9839 | 0.9849 | 0.9491 | 0.9291 | 0.9397 | 0.9390 | 0.8795 |
| | random 5-mer | 768 | 18310562 | **0.9860** | 0.9866 | 0.9513 | 0.9426 | 0.9472 | 0.9470 | **0.8945** |

**Table 2.** Comparison of different k-mer embeddings used in the Deepromoter model. The model parameters are selected accroding to the MCC performance on the development set. The best AUROC and MCC scores are shown in bold font.

### 4. Experiments supplements

#### 4.1 Other evaluation metrics on TBFS tasks

The performance (AUROC, AUPRC, F1, and MCC) of each ChIP-seq dataset is provided in the corresponding task fold. Besides the AUROC shown in the main text, the overall boxplots of other evaluation metrics are provided in the following.

##### 4.1.1 TBFS motif discovery (690)

##### AUPRC

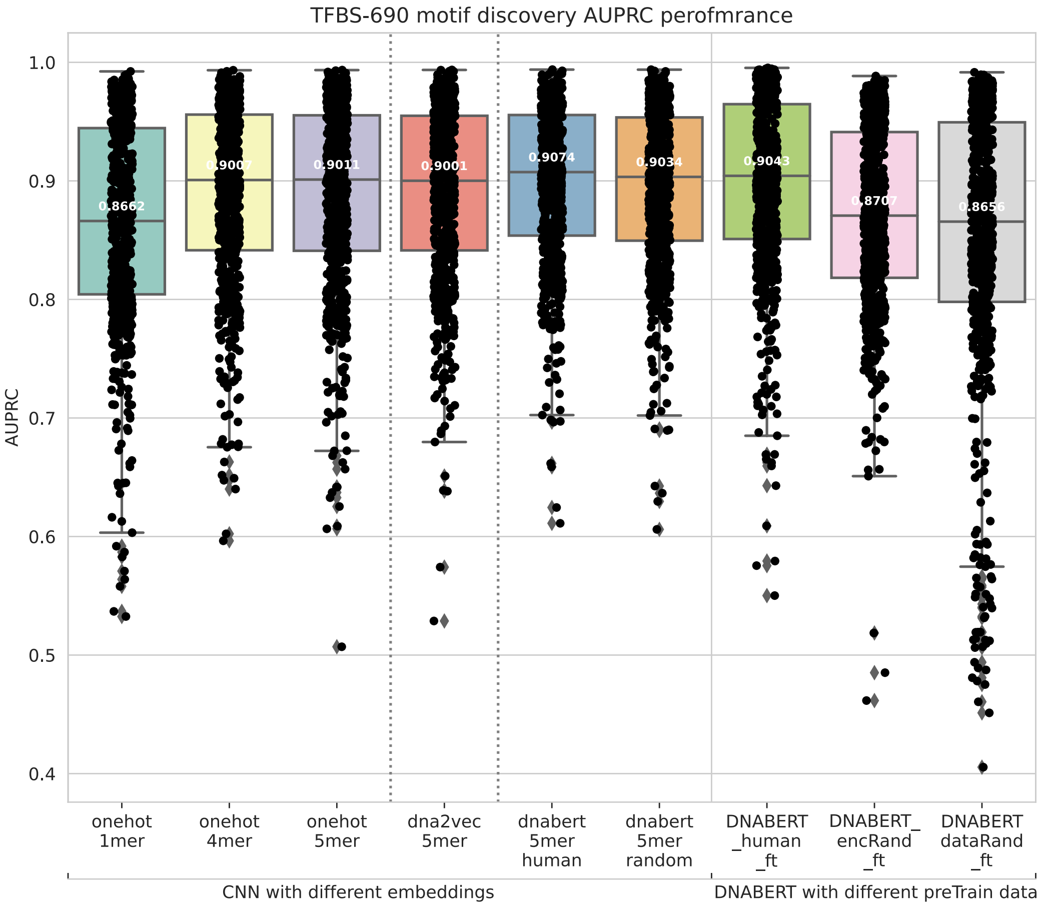

**F1**

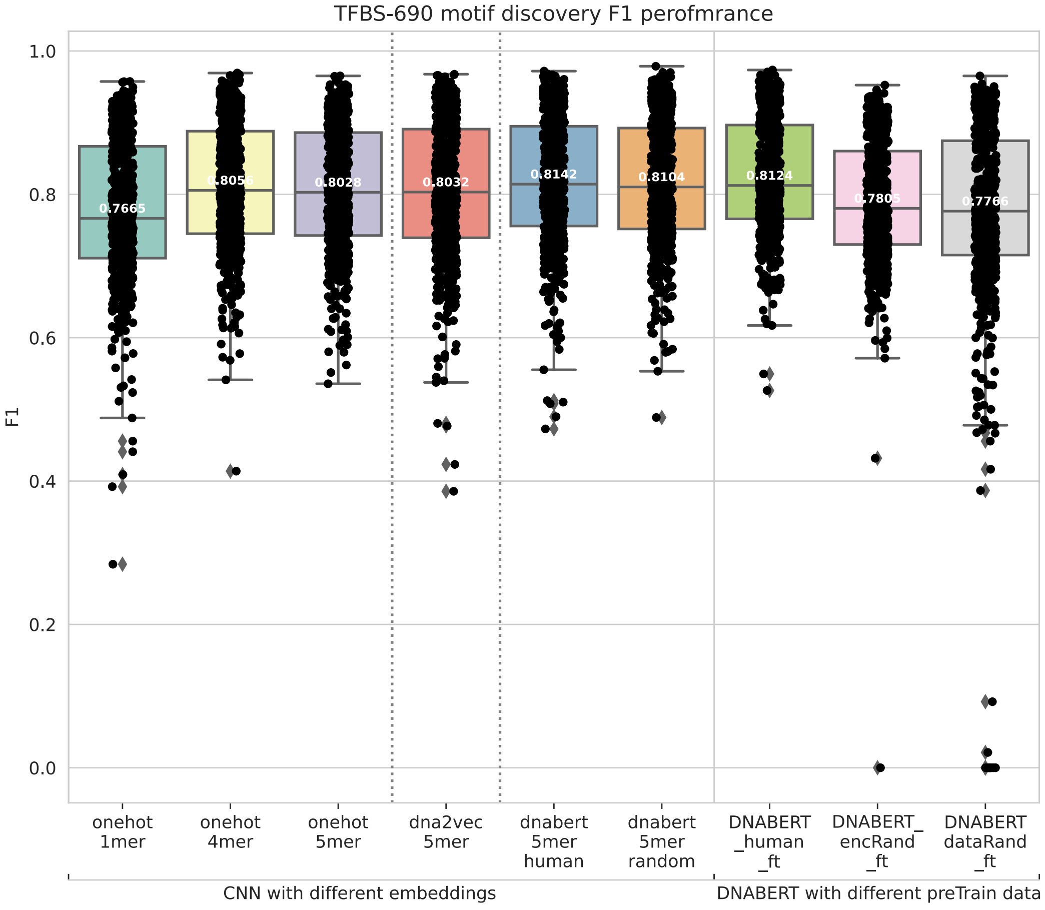

TFBS-690 motif discovery F1 perofmrance
F1
1.0
0.8
0.6
0.4
0.2
0.0
0.7665
0.8056
0.8028
0.8032
0.8142
0.8104
0.8124
0.7805
0.7765
onehot 1mer
onehot 4mer
onehot 5mer
dna2vec 5mer
dnabert 5mer human
dnabert 5mer random
DNABERT _human _ft
DNABERT_ encRand _ft
DNABERT dataRand _ft
CNN with different embeddings
DNABERT with different preTrain data

**MCC**

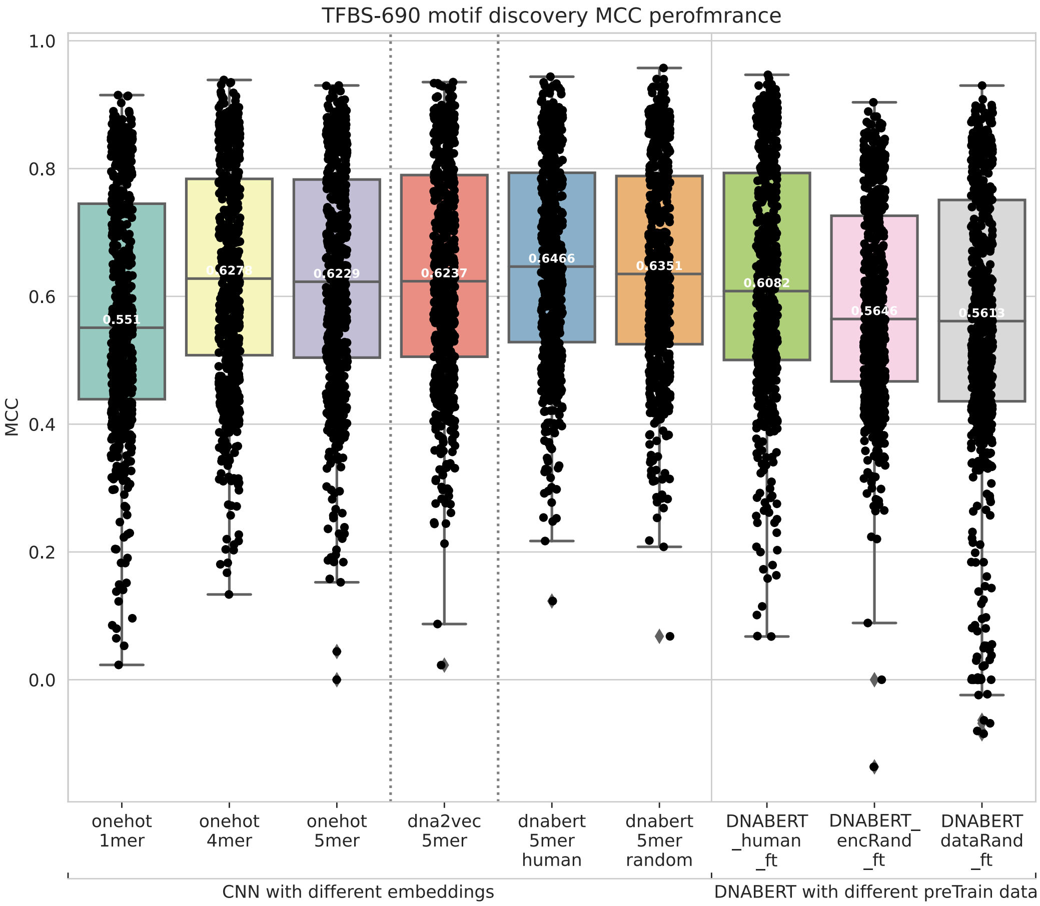

TFBS-690 motif discovery MCC perofmrance
MCC
1.0
0.8
0.6
0.4
0.2
0.0
0.551
0.6278
0.6229
0.6237
0.6466
0.6351
0.6082
0.5646
0.5613
onehot 1mer
onehot 4mer
onehot 5mer
dna2vec 5mer
dnabert 5mer human
dnabert 5mer random
DNABERT _human _ft
DNABERT_ encRand _ft
DNABERT dataRand _ft
CNN with different embeddings
DNABERT with different preTrain data

##### 4.1.2 TBFS motif occupancy identification (420)

##### AUPRC

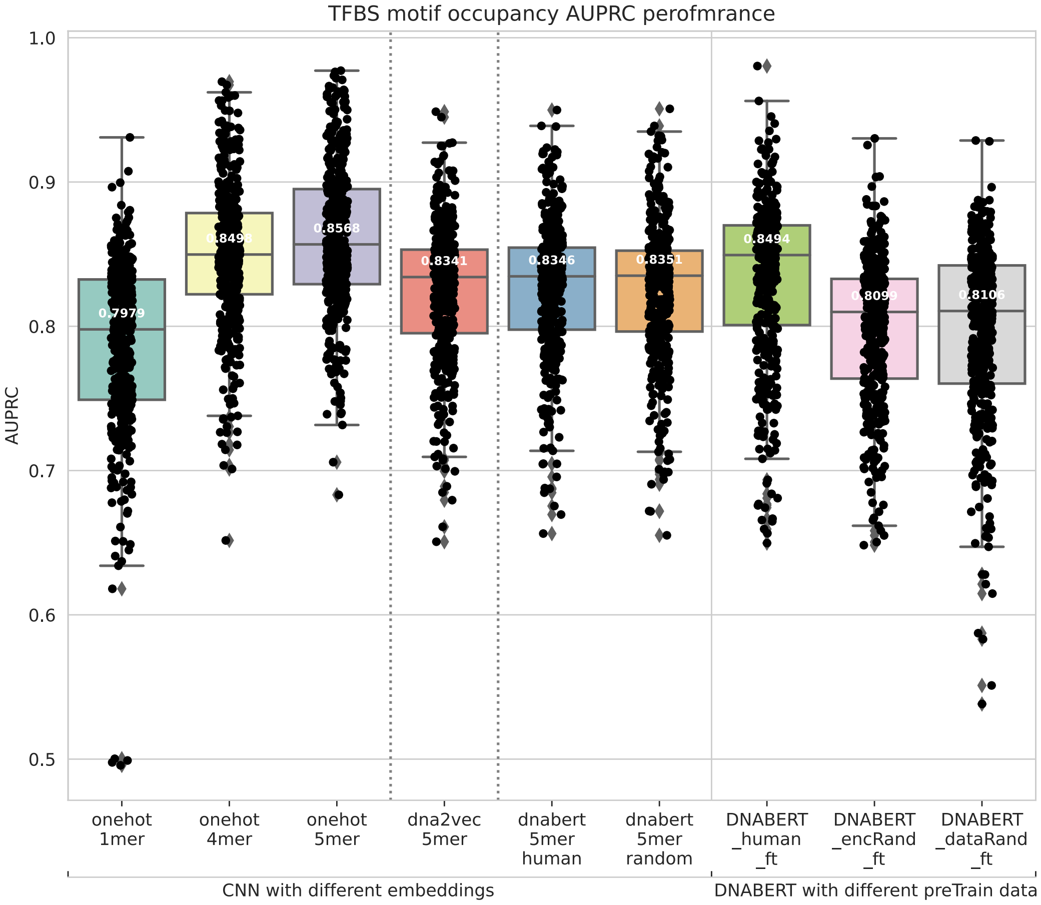

# F1

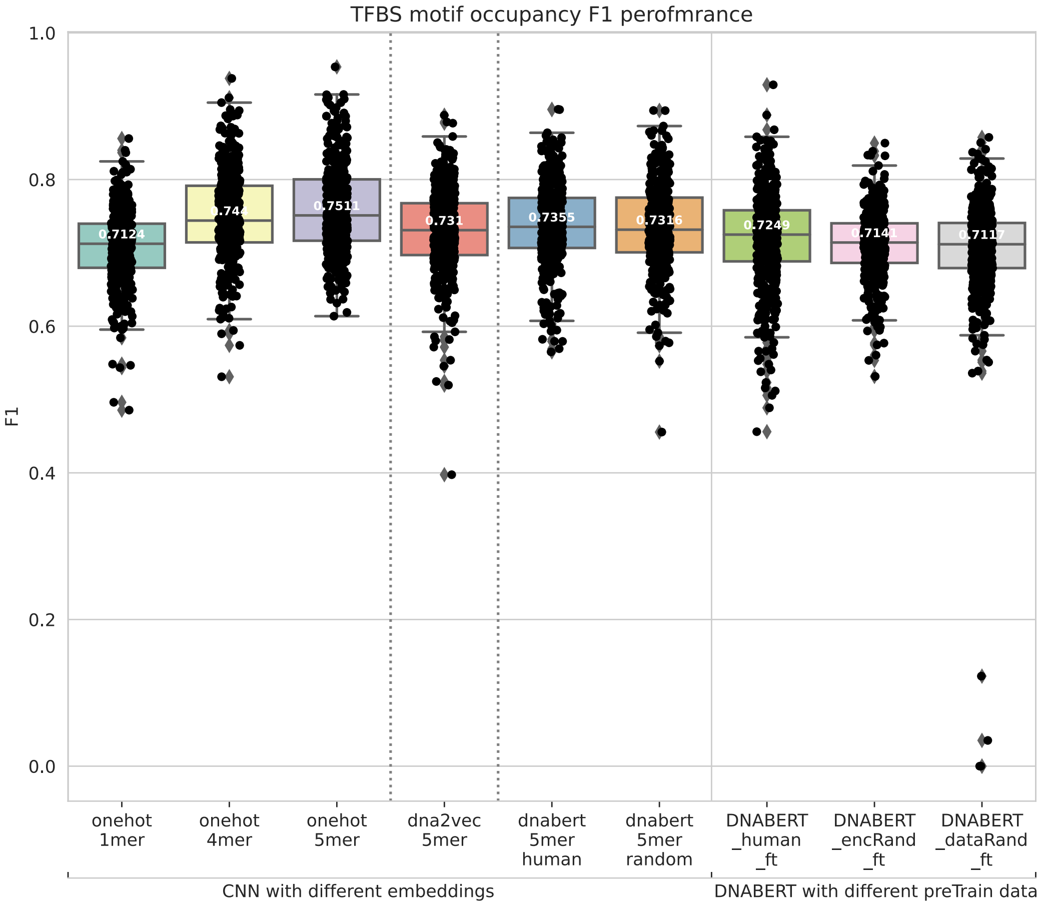
TFBS motif occupancy F1 perofmrance
F1
1.0
0.8
0.6
0.4
0.2
0.0
0.7124
0.7511
0.731
0.7355
0.7316
0.7249
onehot 1mer
onehot 4mer
onehot 5mer
dna2vec 5mer
dnabert 5mer human
dnabert 5mer random
DNABERT _human _ft
DNABERT _encRand _ft
DNABERT _dataRand _ft
CNN with different embeddings
DNABERT with different preTrain data

### MCC

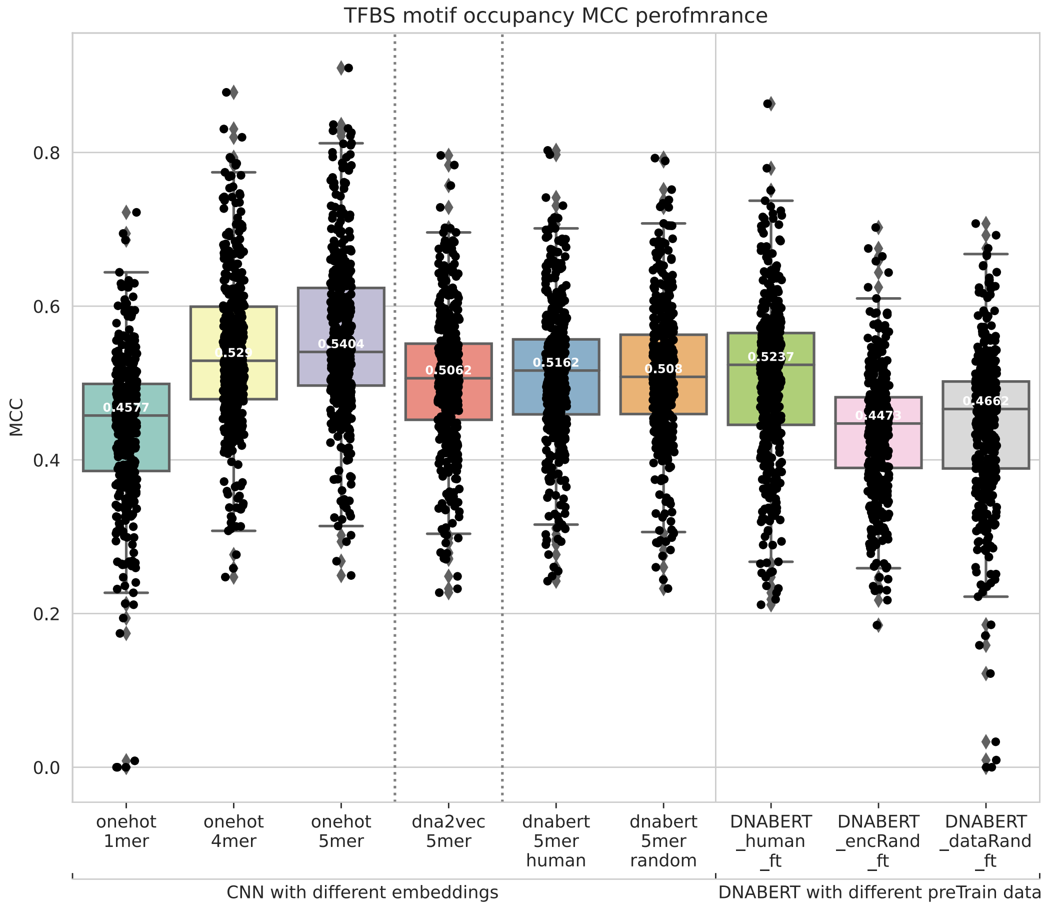

TFBS motif occupancy MCC perofmrance
MCC
0.0
0.2
0.4
0.6
0.8
0.4577
0.529
0.5404
0.5062
0.5162
0.508
0.5237
0.4473
0.4662
onehot 1mer
onehot 4mer
onehot 5mer
dna2vec 5mer
dnabert 5mer human
dnabert 5mer random
DNABERT _human _ft
DNABERT _encRand _ft
DNABERT _dataRand _ft
CNN with different embeddings
DNABERT with different preTrain data
